# Integrating carbon utilization and transport processes into a crop growth model enables the prediction of emergent soybean carbon allocation behavior

**DOI:** 10.64898/2026.08.27.747615

**Authors:** Ximin Piao, Edward B. Lochocki, Justin M. McGrath, Megan L. Matthews

## Abstract

Accurately modeling carbon (C) allocation is essential for predicting crop yield and the performance of new cultivars in various environments. Most crop models allocate C empirically, using fixed partitioning tables or harvest indices that prescribe allocation without representing the underlying physiology, limiting their predictive power under novel conditions. A mechanistic alternative, in which C allocation emerges from local utilization and transport, could instead respond dynamically to environmental changes, source-sink perturbations, and organ-level trait modifications. To achieve this design, we integrated a utilization-transport-resistance (UTR) allocation model into the Soybean-BioCro crop growth modeling framework. We calibrated and validated the model using organ biomass data from two soybean cultivars grown at two CO_2_ levels over eight seasons, achieving accuracy comparable to partitioning-based models while predicting more reasonable carbon allocation fractions. Further, the UTR-BioCro model predicted leaf and stem total nonstructural carbohydrate concentrations with reasonable accuracy compared to experimental measurements across the 2022 growing season. A local sensitivity analysis of the model parameters indicated that the onset of reproductive growth influenced yield more strongly than utilization or transport parameters suggesting the timing of this transition as a potential target for crop improvement. Finally, the UTR-BioCro model reproduced yield responses to source-sink perturbations including shading and pod removal, and captured the qualitative response to defoliation without requiring scenario-specific tuning as most partitioning approaches require. By grounding C allocation in physiological mechanisms, this work provides a foundation for predicting crop responses across diverse environments and engineered traits, supporting crop improvement for a changing environment.

## 1 Introduction

Soybean is the primary source of plant-based protein in animal feed and human diets (Qin et al., 2022), and global demand is projected to rise by 80% between 2005–2007 and 2050 (Fischer et al., 2014). Meeting this demand while limiting agricultural expansion requires sustained yield gains. Yet yield decreases are projected across major growing regions by the end of the century without adaptation (Challinor et al., 2014; Hasegawa et al., 2022). Biotechnology and targeted breeding are increasingly seen as essential tools to accelerate adaptation, but their design depends on a systems-level understanding of how molecular and metabolic interventions propagate to whole-plant growth (Rezaei et al., 2023). Crop models that integrate physiological and environmental responses can support this design process, provided their underlying mechanisms respond to novel conditions outside the calibration data (Piao & Matthews, 2026).

A key limitation of current crop models is the reliance on empirical relationships. The variation in these empirical relationships contributes to large variability in yield projections under future climates (Dhakar et al., 2018; Heinicke et al., 2022; Jägermeyr et al., 2021; Kothari et al., 2022; Müller et al., 2021). One critical component that lacks physiological representation is the allocation of photoassimilated carbon (C) among plant organs (Camargo-Alvarez et al., 2023; Hartmann et al., 2020). C allocation mediates trade-offs to obtain different resources. Greater allocation to roots enhances drought tolerance and nutrient acquisition but constrains aboveground growth, whereas greater allocation to leaves can either increase or decrease canopy photosynthesis depending on the extent of self-shading, while also increasing respiratory costs as the canopy matures. Shifting allocation toward reproductive organs was central to the Green Revolution, raising yield while reducing lodging risk (Khush, 2001). Given its central role in plant development, accurately modeling how C is dynamically allocated is critical to improving the accuracy of crop model predictions.

In crop models, carbon allocation is typically represented using one of three empirical approaches: a partitioning model, a teleonomic model, or a sink competition model. Partitioning models, or “descriptive allometry models” (Marcelis et al., 1998), prescribe the fraction of photoassimilated C that is allocated to each organ for each development stage (Boote et al., 1998; Hoogenboom et al., 2019; Steduto et al., 2009), sometimes with stress-specific adjustments (Camargo-Alvarez et al., 2023; Enders et al., 2023). Because these fractions vary with environment and cultivar (Ainsworth et al., 2002; Hossain & Li, 2021; Li et al., 2024; Suwa et al., 2010), partitioning models typically require re-calibration for each environmental condition. Teleonomic models assume C is allocated in a way that optimizes for a certain goal, e.g. maximizing biomass or canopy assimilation, or balances shoot photosynthetic capacity against root nutrient uptake capacity to achieve a “functional balance” in acquiring these resources (Johnson & Thornley, 1985; Le Roux et al., 2001; West, 1993). Another form of teleonomic model is the allometric model which assumes power-law relationships among organs following the functional balance principle (Le Roux et al., 2001; West et al., 1997; West, 1993). These approaches presume a centralized “decision-making” mechanism (Brown et al., 2014), which is widely applied but also debated (McConnaughay & Coleman, 1999; Robinson, 2023; Thornley, 1995). Sink competition models treat sinks as competitors for C according to their predefined maximum growth potentials (Coussement et al., 2020; Heuvelink, 1996; Holzworth et al., 2014), but these maxima are typically defined empirically and often do not reflect extreme conditions (Minchin & Lacointe, 2005).

A more mechanistic model for C allocation has been developed based on C utilization and transport processes (Thornley, 1972). Thornley (1972) developed this utilization-transport-resistance (UTR) framework to simulate steady-state vegetative growth, showing that partitioning patterns shift with the assimilation rate and model parameters related to utilization and transport. Variations of the original UTR model have been developed to examine dry-matter partitioning response to root temperature changes with a tomato shoot-root model (Cooper & Thornley, 1976), extending the framework to five organ compartments in a stand-level forest model (Thornley, 1991), adding phosphorate substrate to the shoot-root model and comparing it with teleomic models (Thornley, 1995), and a two-state model with substrate and structural C pools to demonstrate maintenance respiration can be an emergent property of substrate use instead of a distinct process (Thornley, 2011). This UTR model separates the C pool of each organ into substrate and structural C, where substrate C comprises of labile metabolites such as glucose, sucrose, and starch, which are often collectively referred to as total nonstructural carbohydrates (TNC). Structural C represents the carbon integrated into cell walls, proteins, and lipids (Chiariello et al., 2000). The framework has since been applied to forest simulation to improve predictions of carbon fluxes in land surface models (Jones et al., 2020; Thornley, 1991), the post-véraison stage in a grape functional structural plant model (Zhu et al., 2019, 2021), and the vegetative growth of tobacco (Wann & Raper, 1984), but has not been applied to simulate allocation at the organ level for field-grown crops for the full life cycle.

Embedding the UTR framework within a crop model could improve the accuracy of crop model predictions under novel scenarios, enhancing the ability of these models to identify target traits that improve yields across a variety of conditions. Since allocation would emerge from local substrate utilization and transport processes in the UTR model rather than from prescribed fractions, a single parameter set could respond directly to environmental and physiological changes. This suggests that environmental changes that impact source and/or sink activities such as elevated CO_2_ concentration, shading, or biomass perturbations such as pod removal, defoliation, and hail damage (Morgan et al., 2005; Parvej et al., 2025; Proulx & Naeve, 2009) would result in altered carbon allocation dynamics without scenario-specific stress modules. Finally, by simulating substrate carbon dynamics at the organ level, the framework would provide a tool for evaluating crop engineering interventions, projecting the whole-plant consequences of organ-specific metabolic modifications such as enhanced lipid biosynthesis in bioenergy crops (Chen et al., 2026; Clark & Schwender, 2022; Dong et al., 2025) or altered transporter kinetics (Ludewig & Flügge, 2013).

In this study, we integrated the UTR model into the Soybean-BioCro (Lochocki et al., 2022; Matthews et al., 2022) crop growth modeling framework. The BioCro framework is set up to work with adaptive ODE solvers from the Boost library (*Boost 1.89.0* 2026) allowing for the stiff equations in the UTR framework to be solved. The modularity of BioCro also allows the previous partitioning modules to be easily swapped with a new set of allocation modules without needing to adjust the rest of the model’s code (e.g. canopy photosynthesis, soil water dynamics). To simulate soybean growth, we extended Thornley’s original UTR framework to include reproductive development and senescence, by adding a transport process to pod that is switched on during reproductive development and converting portions of the substrate and structural C pools to litter for each tissue during senescence (Figure 1). We calibrated this UTR-BioCro model following the same procedure as the original Soybean-BioCro model (Matthews et al., 2022), where model parameters were fit to biomass data from one soybean cultivar grown over the 2002 and 2005 growing seasons under ambient CO_2_. We validated the model predictions against organ biomass data from the same cultivar grown across two CO_2_ levels during the 2002, 2004-2006 growing seasons and from a different cultivar grown during the 2021-2024 growing seasons. We further validated the UTR-BioCro model’s ability to simulate substrate carbon dynamics with leaf and stem TNC concentrations measured during the 2022 growing season. Next, we used a local sensitivity analysis to identify the parameters that had the largest impact on final yield predictions. Finally, we evaluated the model’s capacity to predict yield responses to source–sink manipulations, including shading, pod removal, defoliation, and hail damage, without scenario-specific tuning of parameters.

**Figure 1:**
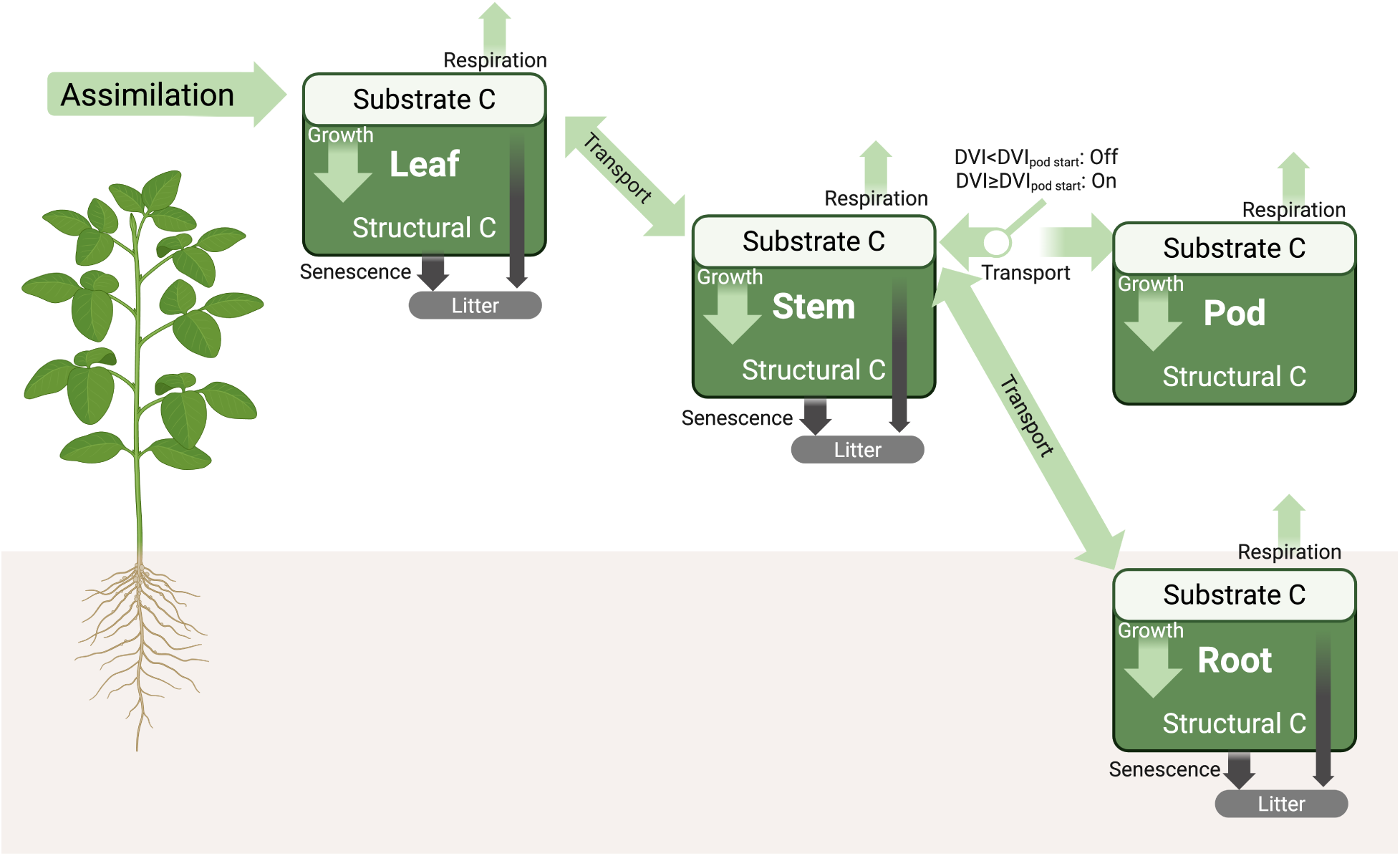
The Utilization-Transport-Resistance (UTR) carbon allocation model adapted from Thornley, 1972 to include reproductive development and senescence.

## 2 Methods

### 2.1 Model Details

#### 2.1.1 The original UTR model

Thornley (1972) represented a vegetative plant as three connected organs, leaf, stem, and root, each with its own pool of substrate C. Two processes govern how the substrate C is distributed. Within each organ, substrate C, *S_O_*, is utilized for growth at a rate, 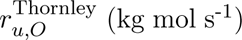, following Michaelis-Menten kinetics (Eq 1),

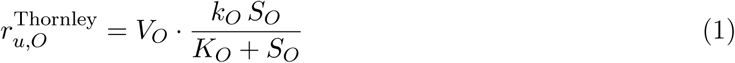

where *O* denotes the organ, *V_O_*is organ volume, *k_O_*the maximum utilization rate, and *K_O_*the substrate concentration at half-maximal rate. A constant fraction of the utilized substrate C, *k_res,O_*, is lost to growth respiration; the remainder is converted to structural mass. Substrate C also moves between connected organs following its concentration gradient (Eq 2),

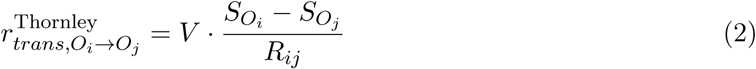

where *R_ij_* is the transport resistance. The carbon transport rate, 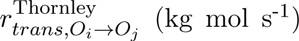, is scaled by the total plant volume *V*. Together with the incoming photosynthate, *P_g_*, utilization and transport define the rate of change of substrate mass for each organ (e.g., for leaf, 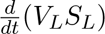 = *P_g_ − r_trans,L→S_ −r_u,L_*), which Thornley (1972) solved for steady-state exponential growth (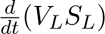 = 0). C transport between organs emerges from the organs’ utilization and transport rates; therefore, unlike the partitioning model, C allocation between organs is not prescribed and will change in response to factors that affect source, sink, or transport relationships. We kept the two essential processes, utilization and transport, but made adaptations so that the model could run at an hourly timestep dynamically with the Soybean-BioCro framework (Lochocki et al., 2022; Matthews et al., 2022) and represent reproductive development. Our adaptations are introduced in the relevant process below.

#### 2.1.2 Utilization and Growth

We replaced Thornley’s Michaelis-Menten utilization (Eq 1) with a Hill equation (Eq 3). Glucose utilization begins with phosphorylation by hexokinase, the rate-limiting enzyme of the first step of glycolysis, which exhibits cooperative substrate binding (Matschinsky, 1996). The resulting sigmoidal response, which the Michaelis-Menten equation cannot capture, produces near-zero utilization at low substrate C and a sharper transition into saturation at high substrate C. The organ substrate C utilization rate, *r_u,O_* (mol m^-2^ hr^-1^), is given by

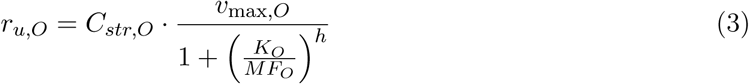

where *O* denotes organ (leaf, stem, root, and pod). *C_str,O_* represents the amount of structural carbon in the organ per land area (mol m*^−^*^2^ ground area). *v*_max_*_,O_* is the maximum utilization rate (hr^-1^). The mass fraction, *MF_O_*, is the ratio of substrate C to structural C (dimensionless); it serves as a proxy for substrate concentration, replacing the volumetric substrate C concentration, *S_O_*, under the assumption that the structural C per land area of an organ, *C_str,O_* (mol m^-2^ ground area), is proportional to the organ volume, *V_O_*. *K_O_* is the dissociation constant of the organ’s utilization rate (dimensionless), representing the *MF_O_* at which the utilization rate is half of *v*_max_*_,O_*. *h* is the Hill coefficient, representing the level of binding cooperativity. We set *h* = 2, within the experimentally reported range of 1.4 to 2.3 for hexokinase (Matschinsky, 1996; Ureta, 1976; Van Schaftingen, 2013). As in Thornley’s model, a fixed fraction, *k_res,O_*, of the utilized substrate C is respired (Eq 4);

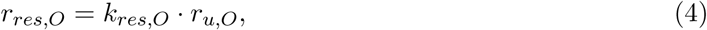

and the remainder becomes structural C (Eq 5),

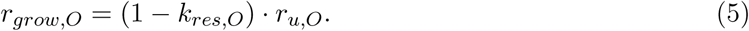

#### 2.1.3 Transport

The transport rate is determined by the mass fraction gradient multiplied by the conductance between the two organs (Eq 6),

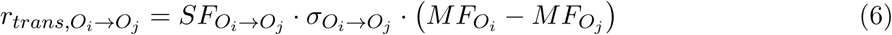

where *O_i_* and *O_j_* represent two connected organs. In the soybean model, the links include leaf to stem, stem to root, and stem to pod. *σ_O__i→Oj_* (hr^-1^) is the conductance of substrate C between the organs. *SF_O__i→Oj_* (mol m^-2^) is the scaling factor accounting for the increasing conductance as the organs grow. Thornley (1972) scaled transport by the total plant volume (Eq 2), however this becomes unrealistic when a new organ begins growing partway through the simulation. In this case, a pod with near-zero mass receives an influx scaled by the whole-plant volume that far exceeds what the pod can utilize. The substrate C concentration then becomes larger in the pod than in the stem, reversing the gradient and causing C to be exported from the pod. To address this limitation, we scaled conductance to the structural C of the smaller of the two organs in each transport link (Eq 7),

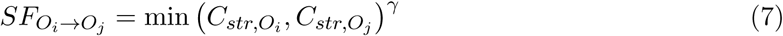

where *γ* is a constant adjusting for the allometric relationship between conductance and structural C. For simplicity, we set *γ* = 1 in the model.

#### 2.1.4 Senescence

The senescence fractions of leaf, stem, and root, *f_sene,O_* (hr^-1^), are determined by the development index (DVI) (Eq 8),

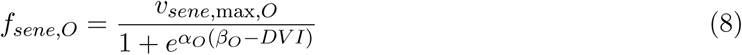

where *v_sene,max,O_* (hr^-1^) is the maximum senescence rate. *α_O_* represents how fast the senescence rate increases as DVI increases. *β_O_*represents the DVI at which the senescence rate has reached half of *v_sene,max_*.

All structural C in the senesced part of the organ becomes litter (Eq 9).

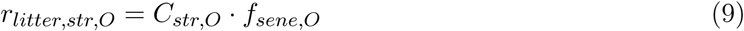

Part of the substrate C in the senesced part of the organ, *r_retain,O_* (mol m^-2^ hr^-1^), is retained in the organ’s substrate C pool (Eq 10), while the rest of the substrate C is assumed to become litter (Eq 11),

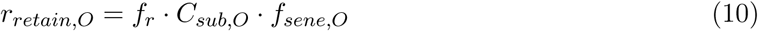

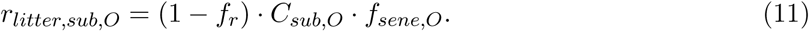

where *f_r_*is the fraction of substrate C retained from senescing part of the organ. The retained substrate C is treated the same as the remaining organ substrate C, which is available for utilization or transport, rather than being remobilized to other organs immediately as in the partitioning model (Matthews et al., 2022). The rate of litter production, *r_litter,O_* (mol m^-2^ hr^-1^), is the sum of senesced structural C and substrate C (Eq 12),

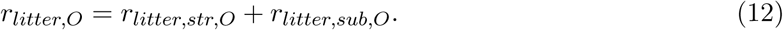

#### 2.1.5 Mass Balance

The change in substrate C (mol m^-2^ hr^-1^) is determined by the C influx and efflux (Eq 13),

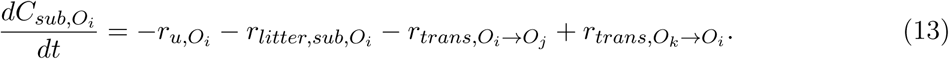

In the leaf, the net assimilation rate, *A_net_*, is also included as a C influx in Eq 13.

The change in structural C (mol m^-2^ hr^-1^) is determined by the growth rate and senescence rate (Eq 14).

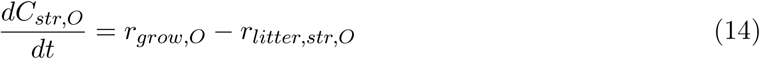

The dry biomass of the organ, *M_O_* (Mg ha^-1^), is calculated as the sum of substrate C and structural C, multiplied by the conversion factor *f* (Eq 15),

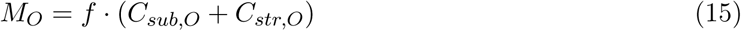

where *f* converts the molar mass (mol m^-2^) to biomass mass (Mg ha^-1^). Assuming the majority of the crop consists of carbohydrates, CH_2_O, we used a conversion factor of 0.3 Mg dry mass ha^-1^ per mol C m^-2^.

#### 2.1.6 Development stage control

Thornley’s model represented only vegetative growth. We extended it to reproductive development by adding a pod organ, whose utilization and transport are activated once the plant enters the early reproductive phase (Figure 1). The development index (DVI) accumulates at a rate determined by thermal time and photoperiod, using the same phenology models as the original Soybean-BioCro (Matthews et al., 2022). Pod utilization and transport processes are activated only after DVI exceeds a threshold corresponding to the onset of stage R3 (beginning pod). This threshold, DVI_pod start_, is treated as a free parameter and estimated by optimization alongside the other model parameters (Table S1). All organ utilization and transport processes cease once the crop reaches physiological maturity, defined by another DVI threshold, DVI_stop growth_ which is similarly calibrated with the other model parameters.

### 2.2 Model parameterization

The parameterization for the UTR model follows the method used to train the original Soybean-BioCro using weighted sum of squared errors (Matthews et al., 2022). The calibrated parameters include (i) maximum utilization rates, *v*_max_*_,O_*, (ii) dissociation constants for the utilization rate, *K_O_*, (iii) respiration fractions, *k_res,O_*, (iv) substrate conductance between connected organs, *σ_Oi→Oj_*, (v) maximum senescence rates, *v_sene,max,O_*, (vi) senescence constants *α_O_* and *β_O_*, (vii) retained fraction of substrate C, *f_r_*, and (viii) the DVI thresholds for the start of pod growth and end of crop growth, DVI_pod start_, DVI_stop growth_ (Table S2). The bounds for each parameter and their final fitted values are summarized in Table S1.

We trained and validated the UTR model using measured organ biomass from 12 cultivar *×* year *×* CO_2_ combinations (Pioneer 93B15 under ambient and elevated CO_2_ for 2002 and 2004–2006, and LD11-2170 under ambient CO_2_ for 2021–2024). Measurements from the soybean cultivar Pioneer 93B15, that were planted at the SoyFACE facility in Urbana, IL, at ambient CO_2_ levels in 2002 and 2005 were used for parameter fitting, while the other years and CO_2_ levels were used for validation (Bernacchi et al., 2005; Bishop et al., 2015; Morgan et al., 2005). In this dataset, the root biomass was not measured directly, so it was inferred from the biomass of the other organs. We also collected biomass and gas exchange measurements from a more recent cultivar, LD11-2170, which was grown at the Energy Farm in Urbana, IL, from 2021 to 2024 under ambient CO_2_ for additional validation. The weather data at SoyFACE came from the SURFRAD Bondville, IL site (Augustine et al., 2005), including hourly radiation, temperature, relative humidity, and wind speed. The precipitation data came from the WARM Champaign, IL dataset (Water and Atmospheric Resources Monitoring Program, 2015) as described in (Matthews et al., 2022). The weather data at the Energy Farm was collected at the field site with a Campbell Scientific ClimaVue 50 sensor mounted at a height of 10 m.

Photosynthetic CO_2_ response curves were measured on LD11-2170 using a portable photosynthesis instrument (LI-6800,LI-COR Biosciences, Lincoln, Nebraska) on Aug 5 in 2022. Parameters of the Farquhar-von-Caemmerer-Berry (FvCB) photosynthetic model were estimated from the curves using maximum likelihood (PhotoGEA v1.3.3; Lochocki et al., 2025). These inferred photosynthetic parameters (Table S3) were used for the LD11-2170 simulations.

### 2.3 Sensitivity Analysis

To identify which fitted parameters most strongly influence final yield, we performed a local sensitivity analysis on the calibrated UTR model parameters (Table S1). Each parameter was individually changed by ±10% of its fitted value while all other parameters were held constant. Sensitivity was quantified as the percent change in final pod mass relative to the baseline. The analysis was repeated across all 12 cultivar *×* year *×* CO_2_ combinations.

### 2.4 Carbon allocation fractions calculation

Because the UTR model does not prescribe allocation fractions, we computed analogous quantities for comparison with the partitioning model using two approaches. The first compares daily carbon usage in each organ to the total daily usage across all organs (Eq 16).

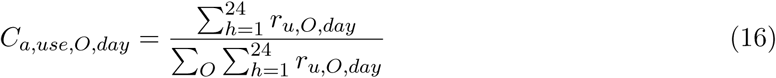

The second approach compares daily net organ imports to the daily net assimilation rates. The rates are aggregated daily to reduce the influence of the large diurnal variability causing uninterpretable results. The allocation fraction for each organ equals its summed daily net flux divided by the daily total carbon input (Eqs 17–20).

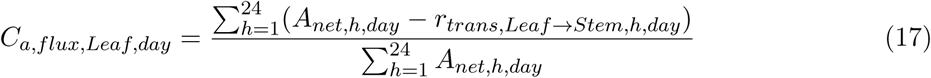

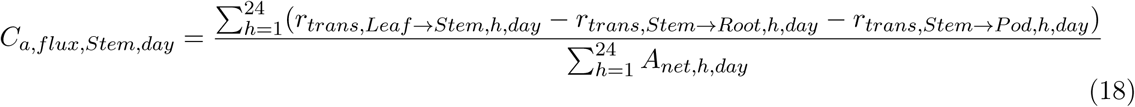

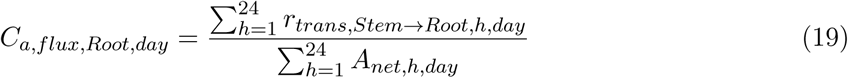

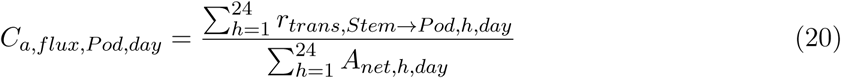

### 2.5 Carbohydrate concentration data

We sampled leaves and stems of LD11-2170 grown at the Energy Farm to measure substrate C concentrations on six days during the 2022 growing season: July 5-6, and July 28 (vegetative phase), August 24, September 15, and October 5 (reproductive phase). On July 5–6, samples were collected at nine time points spanning a 24-hour cycle at 3-hour intervals. On October 5, only one stem sample was taken around noon since all leaves had fallen. On the other dates, samples were collected around dawn, noon, and dusk. At each time point, we sampled four plants, taking 1.04 cm-diameter discs from the top fully expanded leaves and 2–3 cm stem sections from the stems closest to the sampled leaves. All samples were immediately frozen in liquid nitrogen and stored at –80 °C until analysis.

TNC was determined using a continuous enzymatic substrate assay (Rogers et al., 2004). Samples were lyophilized to constant weight to preserve TNC (Chiariello et al., 2000; Eyles et al., 2013). They were then weighed, pulverized in a tissue lyser, and extracted four times with HEPES-buffered ethanol at 80 °C for 20 minutes each. The supernatant was used for soluble sugar analysis (glucose, fructose, and sucrose); the remaining pellet was used for starch determination.

For soluble sugars, samples and glucose standards (40 µL each) were added to a clear assay plate, followed by 160 µL assay buffer (0.1 M HEPES with MgCl_2_, ATP, NADP mix, and G6PDH). Hexokinase, phosphoglucose isomerase (PGI), and invertase (2 µL each) were added sequentially. After each enzyme addition, absorbance at 340 nm was measured on using a microplate spectrophotometer (Synergy HTX and Gen5 software, BioTek) until stabilization, and the maximum optical density was used for calculations. Glucose, fructose, and sucrose concentrations were calculated from a quadratic fit to the glucose standard curve. For starch, the pellet was digested to glucose overnight in 50 mM sodium acetate with amyloglucosidase and *α*-amylase (both MillporeSigma), and the resulting glucose concentration was quantified using the same assay. Measurement precision was assessed as the coefficient of variation (CV) between the replicates; results with CV above 10% were discarded.

### 2.6 Source-Sink Manipulation Scenarios

To evaluate whether the UTR model captures carbon allocation changes under source-sink manipulation, we compared the yield responses predicted by the UTR and partitioning models against published experimental results. We simulated these scenarios for the 2002, 2004-2006 growing seasons in Urbana, IL, using the parameterization for Pioneer 93B15. The field experiments being emulated were performed elsewhere and on different cultivars. The canopy shading and pod removal experiments were conducted at Becker, St. Paul, and Rosemount, MN, in 2006 and 2007 using the cultivar Pioneer 91M61 (Proulx & Naeve, 2009). The defoliation experiment was conducted at Ames, IA, and West Lafayette, IN, from 2016–2018 using five cultivars (Parvej et al., 2025).

Shading was imposed by reducing the “solar” input in the weather file by 50%, 60%, 70%, and 80% after R5 (DVI = 1.5) to reflect the shading intensities in the experiment (Proulx & Naeve, 2009). Pod removal was simulated by reducing the pod mass (substrate and structural C) by 20%, 40%, 60%, and 70% at R5 reflecting the experimental levels of pod removal. The experimental levels of defoliation were simulated by reducing leaf mass by 0%, 25%, 50%, 75%, and 99.9% at R4 and R5. We also simulated a hail event that occurred on DOY 198 in 2003 during the vegetative phase (V7) that caused 60% leaf loss and 21% aboveground biomass loss (Morgan et al., 2005). To simulate the hail event, the substrate and structural C were reduced by 60% in the leaf and 50% in the stem on DOY 198. The 50% stem reduction approximates the combined loss of fallen stem tissue and damaged tissue that remained on the plant but lost physiological functionality.

The DVI values corresponding to stages R4 (full pod) and R5 (beginning seed) (Fehr et al., 1971) were estimated from the biomass observations. For Pioneer 93B15 grown in 2002 and 2004–2006, we examined the simulated DVIs on the days where seed mass was first reported and on the preceding measurement days. The first reported seed measurements had an average DVI of 1.53, ranging 1.42–1.66, and the DVIs of preceding measurements averaged 1.34, ranging 1.24–1.44. Because R5 marks the onset of seed development “at one the four uppermost nodes with a completely unrolled leaf” (Fehr et al., 1971), it was approximated as DVI = 1.5, slightly lower than the average of the first-seed measurements and above the upper bound (1.44) of the preceding measurements. R4 was approximated as DVI = 1.35, close to the average DVI of the preceding measurement, and below the lower bound (1.42) of the first-seed measurements. The DVI corresponding to R4 is also confirmed by calculating the DVIs for the days when the simulated pod mass reaches the final shell mass from the experimental measurements, since shell growth dominates pod growth first and mostly completes by R4 (full pod) by definition (Fehr et al., 1971). The corresponding DVIs are 1.35, 1.3, 1.36, and 1.29 for 2002, 2004–2006, respectively, similar to our estimated DVI of 1.35 for R4.

In the UTR-BioCro model, we estimate the biomass of the entire pod which is composed of both the shell and seed. To compare the predicted changes in yield against the published experimental yield changes we assumed a fixed seed:pod ratio. This assumption is supported by literature for pod removal (Kollman et al., 1974; Schonbeck et al., 1986), canopy shading (Egli & Bruening, 2005; Schou et al., 1978), and defoliation before R5 (Board et al., 2010; Poudel et al., 2025). This assumption may not hold for defoliation at R5 which we discuss in Section 4.2.

## 3 Results

### 3.1 UTR-BioCro predicts biomass across cultivars and CO_2_ levels over 8 years

The UTR-BioCro model was parameterized using leaf, stem, and pod biomass measurements from the Pioneer 93B15 soybean cultivar grown at SoyFACE under ambient CO_2_ (372 ppm) in 2002 and 2005 (Figure 2A&C). The root mean squared errors (RMSEs) between the predicted and measured biomass for these parameterization years were 0.57 and 0.62 Mg ha^-1^ respectively. The predicted biomasses were then validated against measurements from the same cultivar grown under ambient CO_2_ in 2004 and 2006 (Figure 2B&D), and under elevated CO_2_ (550 ppm) in 2002, 2004-2006 (Figure 2E-H). The RMSEs between predicted and measured biomasses were of similar magnitude to those from the parameterization years, ranging from 0.52 to 1.09 Mg ha^-1^.

**Figure 2:**
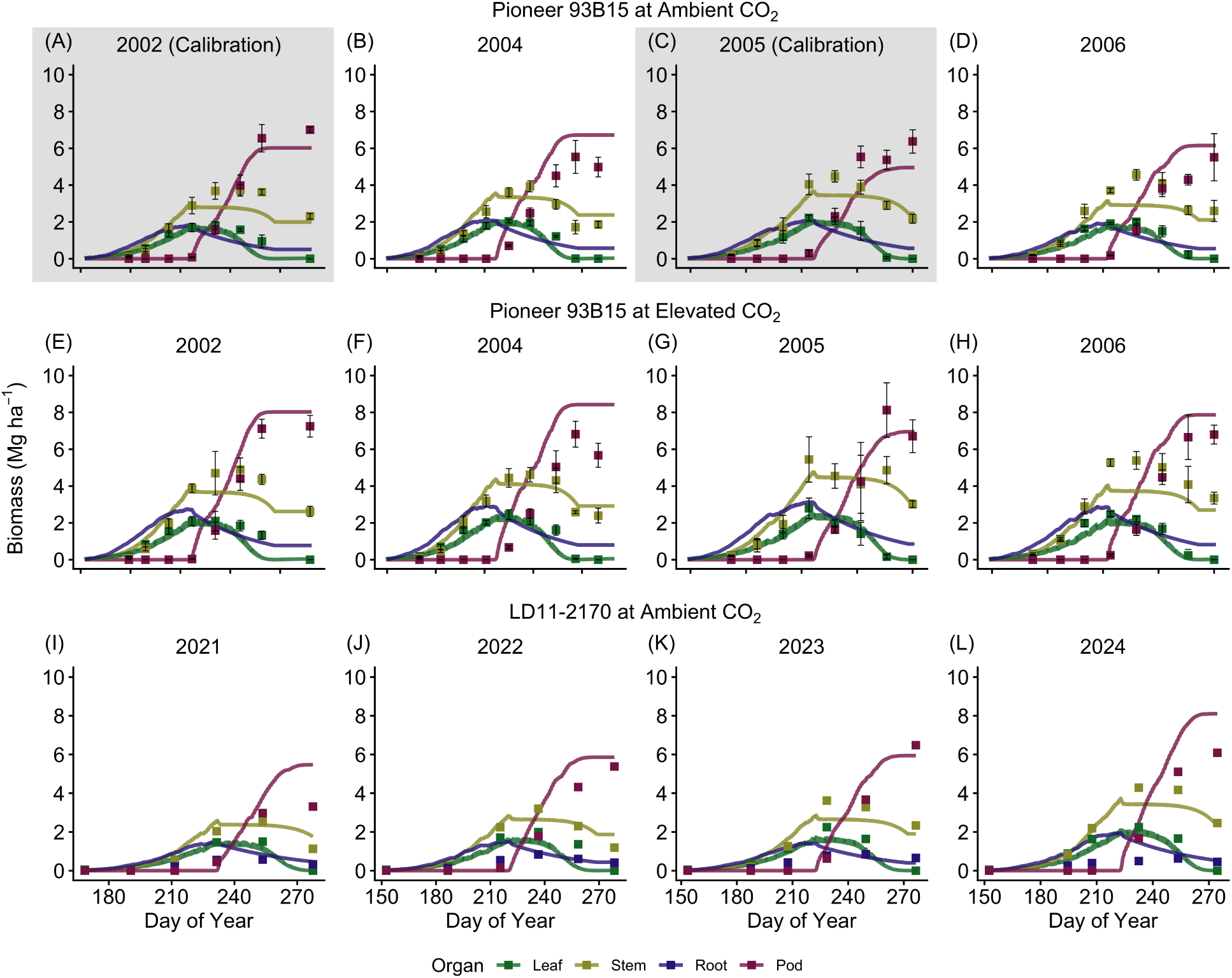
Simulated organ biomasses from the UTR-BioCro model compared with the field measurement. (A-D) Pioneer 93B15 planted at ambient CO_2_. Data from 2002 and 2005 (shaded) were used for model parameterization, (E-H) Pioneer 93B15 grown at elevated CO_2_ concentration, (I-L) LD11-2170 grown at ambient CO_2_ in 2021-2024. Squared symbols (with error bars for Pioneer 93B15) represent measured data (mean *±* SD). Lines represent simulated biomass.

Using the same UTR model parameterization (Table S1), we then simulated a more modern soybean cultivar, LD11-2170 (Table S3), grown under ambient CO_2_ levels (414.70–422.80 ppm in 2021–2024). The RMSEs between the predicted and measured LD11-2170 biomasses were 0.63, 0.61, 0.51, and 0.79 Mg ha^-1^ for 2021 to 2024, respectively (Figure 2I-L).

Overall, the leaf, stem, and pod biomasses predicted by the UTR-BioCro model were consistent with the measured biomasses. The final pod biomasses in 2021 and 2024 were the most different from the measurements (Figure 2 I&L). The field plots where the final biomass measurements were collected in 2021 experienced damage from rabbits which may have resulted in lower than expected experimental pod measurement despite similar leaf, stem, and root biomass measurements and predictions as 2022-2024. The accumulative precipitation during the mid reproductive stage (DVI from 1.2 to 1.8) was the lowest in 2024 (2.78 mm) compared to the other years (86.70 mm, 120.65 mm, and 62.40 mm for 2021 to 2023). Although BioCro’s soil water model adjusts stomatal conductance under water stress and thereby reduces the assimilation rate, the current model does not account for slower growth or accelerated senescence under water stress which could explain the discrepancy for 2024.

### 3.2 Carbon allocation fractions, carbohydrate storage and mobilization, and root:shoot ratio are emergent properties of the UTR-BioCro model

Unlike the partitioning model, which prescribes fixed allocation fractions, these fractions emerge from the system dynamics in the UTR model. For comparability, we used two methods to calculate allocation fractions: (i) organ carbon usage fraction method (Eq 16, Figure 3A), and (ii) daily organ net import method (Eq 17-20, Figure 3B). The two methods yield broadly consistent carbon allocation trends, differing only in the precise allocation percentages. While the general allocation pattern remains consistent across years, daily fractions vary in response to environmental conditions and organ-level metabolic activity.

**Figure 3:**
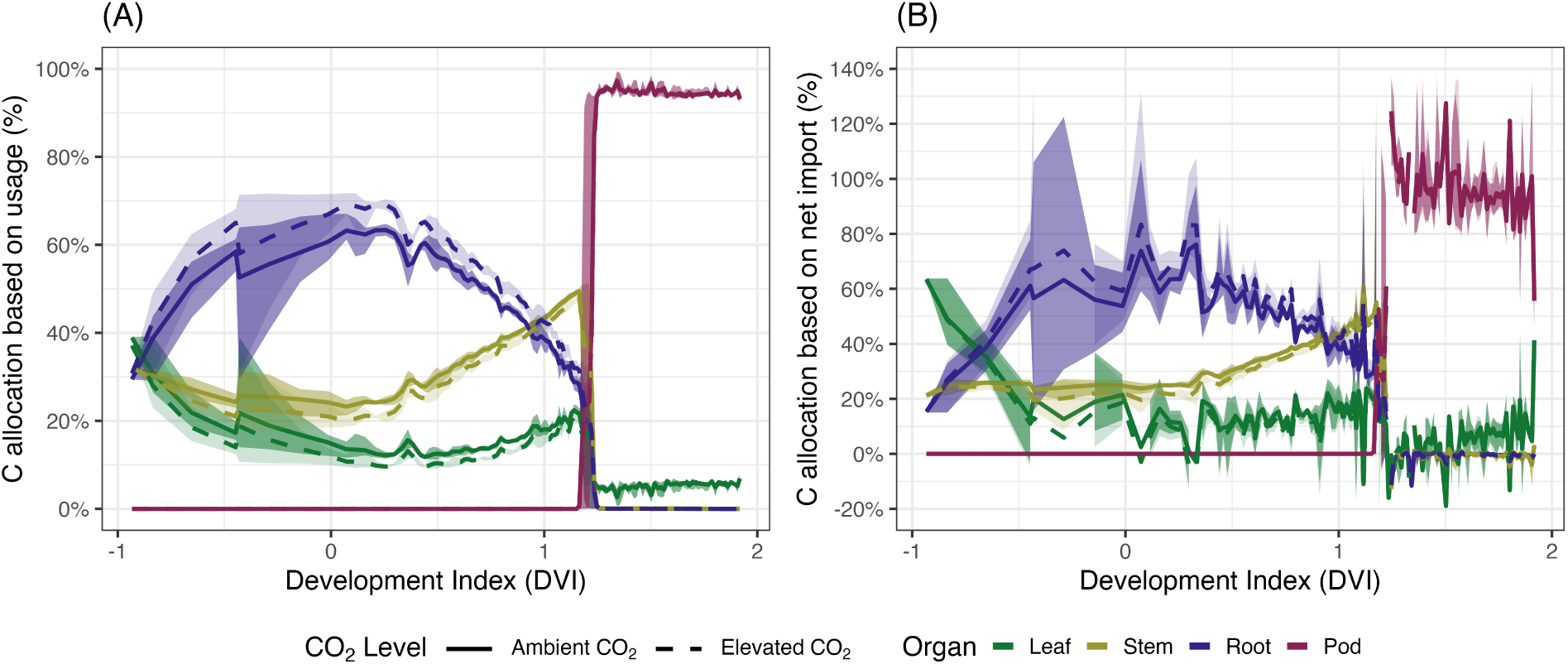
Average daily carbon use fractions of development index (DVI) for Pioneer 93B15 at ambient (solid line) and elevated (dashed line) CO_2_ based on (A) usage and (B) flux. Darker (ambient CO_2_) and lighter (elevated CO_2_) shaded bands indicate the corresponding ranges across years.

Based on the carbon usage fractions (Figure 3A), during the pre-emergence stage (DVI<0), approximately 60% of the net photosynthate is allocated to the roots while about 25% and 15% are allocated to the leaves and stem respectively, consistent with the common observation that the radicle (the embryonic root tissue) dominates growth during the pre-emergence phase (Huang et al., 2014). During the vegetative phase and early reproductive phase (0<DVI<1.2), leaf allocation fluctuates between 10% and 20%, stem allocation rises to approximately 50%, and root allocation decreases to approximately 40%. Once pod growth begins (DVI>1.2), pod usage increases to above 95%. To support this allocation of carbon to the pod, the fraction of carbon usage of the vegetative organs decreased to slightly above 0%.

During the reproductive phases, the net import method calculates that up to 140% of the daily assimilated carbon is allocated to the pod, which is accompanied by near-zero or negative carbon allocation to the vegetative organs (Figure 3B). In contrast, the usage fraction method (Figure 3A) indicates that leaf and stem continue to grow, albeit at a reduced rate, throughout the reproductive phases. This discrepancy arises because the gross canopy assimilation rate does not equal the sum of carbon usage in all organs at each timestep (Figure S4). Instead, assimilate fixed during the vegetative stage is retained in substrate pools and drawn down during the reproductive stage. This delayed carbon usage demonstrates that the UTR model implicitly simulates vegetative-stage carbon storage and its subsequent reproductive-stage mobilization as an emergent property, without imposing these processes.

While the two methods characterize carbon allocation at each timestep, cumulative carbon use reveals where carbon is invested throughout the lifecycle (Figures 4, S1). Leaf, stem, and root grow exponentially during the vegetative stage (Figure 4A). Throughout the growing season, over 50% of assimilated carbon is lost to respiration, with respiration from the leaves, stems, roots, and pods accounting for 47%, 3%, 3%, and 5% of the gross assimilation, respectively, at ambient CO_2_ (Figures 4B, S2).

**Figure 4:**
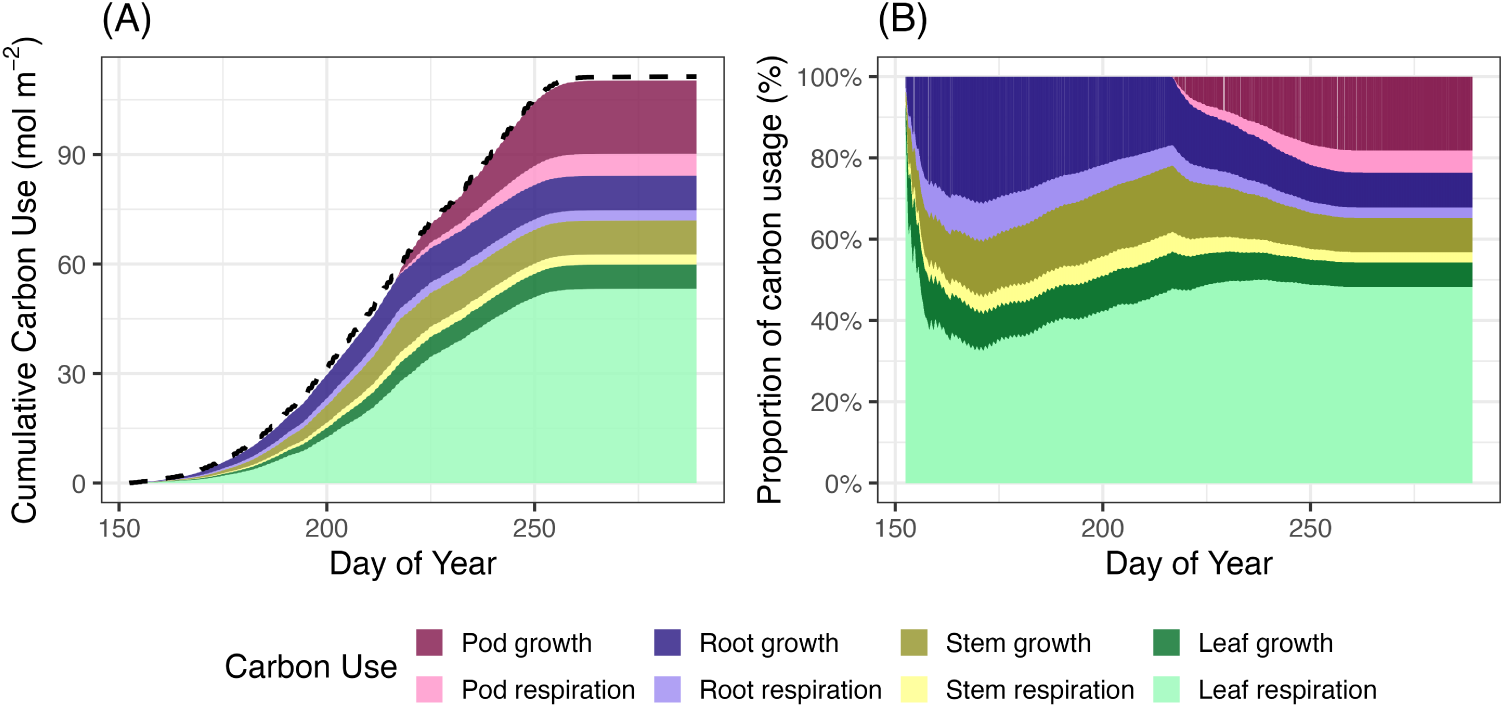
(A) Cumulative carbon use in Pioneer 93B15 at ambient CO_2_ in 2002 from the UTR model. The dashed line represents the cumulative gross canopy assimilation rate. (B) Fractions of organ cumulative carbon use compared to the total use.

An increased root:shoot ratio (R/S) is often observed under elevated CO_2_ (Li et al., 2024; Rogers et al., 1992; Rogers et al., 1995), although the response is inconsistent for soybean (Section 4.3). Both the UTR and partitioning models predict greater biomass under elevated CO_2_, but their R/S predictions diverge: the UTR model predicts a sustained increased in R/S across the season up to 1.4 (Figure 5) whereas the partitioning model predicts an apparent increase only during the early vegetative stage up to 15.6.

**Figure 5:**
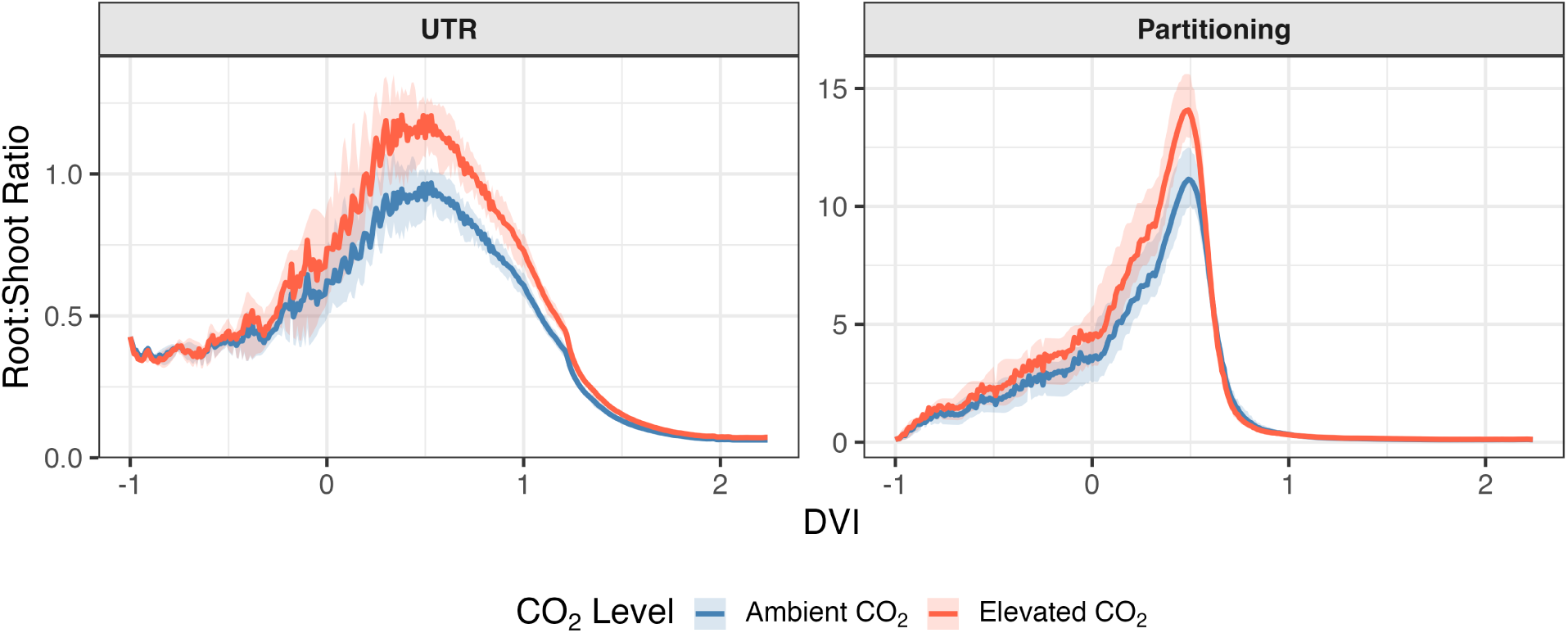
Predicted root:shoot ratios by the UTR and partitioning models for Pioneer 93B15 across the growing season at ambient (blue) and elevated (red) CO_2_. Root:Shoot ratio is the ratio of root biomass to shoot biomass (including leaf, stem, and pod). The lines represent the average ratio for each CO_2_ level across the four years (2002, 2004-2006). The shaded regions represent the range of the predicted ratios across the four years.

Elevated CO_2_ changes the R/S ratios in the two models through different mechanisms. In the UTR model, elevated CO_2_ raises the photosynthetic rate and, consequently, substrate concentrations throughout the plant. Because carbon utilization in above-ground organs approaches saturation at high substrate concentrations (Figure S8), further increases in substrate concentration yield diminishing gains in utilization there. A larger share of photosynthate is therefore exported to below-ground tissue instead (Figure S9), raising the R/S. The partitioning model’s early-stage increase has a different mechanism: elevated CO_2_ boosts canopy assimilation more at the early stage when root allocation is high (nearly 100%). This effect disappears as allocation shifts toward the shoot and the canopy approaches closure.

### 3.3 Parameters specific to the reproductive developmental phases have strongest impact on final yield

The sensitivity of final pod mass to ±10% changes from the fitted parameters values was examined across the 12 cultivar *×* year *×* CO_2_ combinations (Pioneer 93B15 under ambient and elevated CO_2_ for 4 years, and LD11-2170 under ambient CO_2_ for 4 years) (Table 1). The final pod mass was most sensitive to the timing of pod growth and leaf senescence. Pod biomass increased by 18.4–53.8% with an earlier start to pod growth (−10% DVI_pod start_), and decreased by 21.5– 34.6% when pod growth was delayed (+10% DVI_pod start_). Earlier senescence (−10% *β_L_*) reduced pod biomass by 15.2–34.8%, and delayed leaf senescence (+10% *β_L_*) increased pod biomass by 7.8–24.7%. Delaying the DVI threshold for the end of growth rarely increased the final pod mass, indicating growth stops as TNC is depleted in most scenarios. However, early termination (−10% DVI_stop growth_) reduced yield by 2.8–6.1%.

**Table 1:**
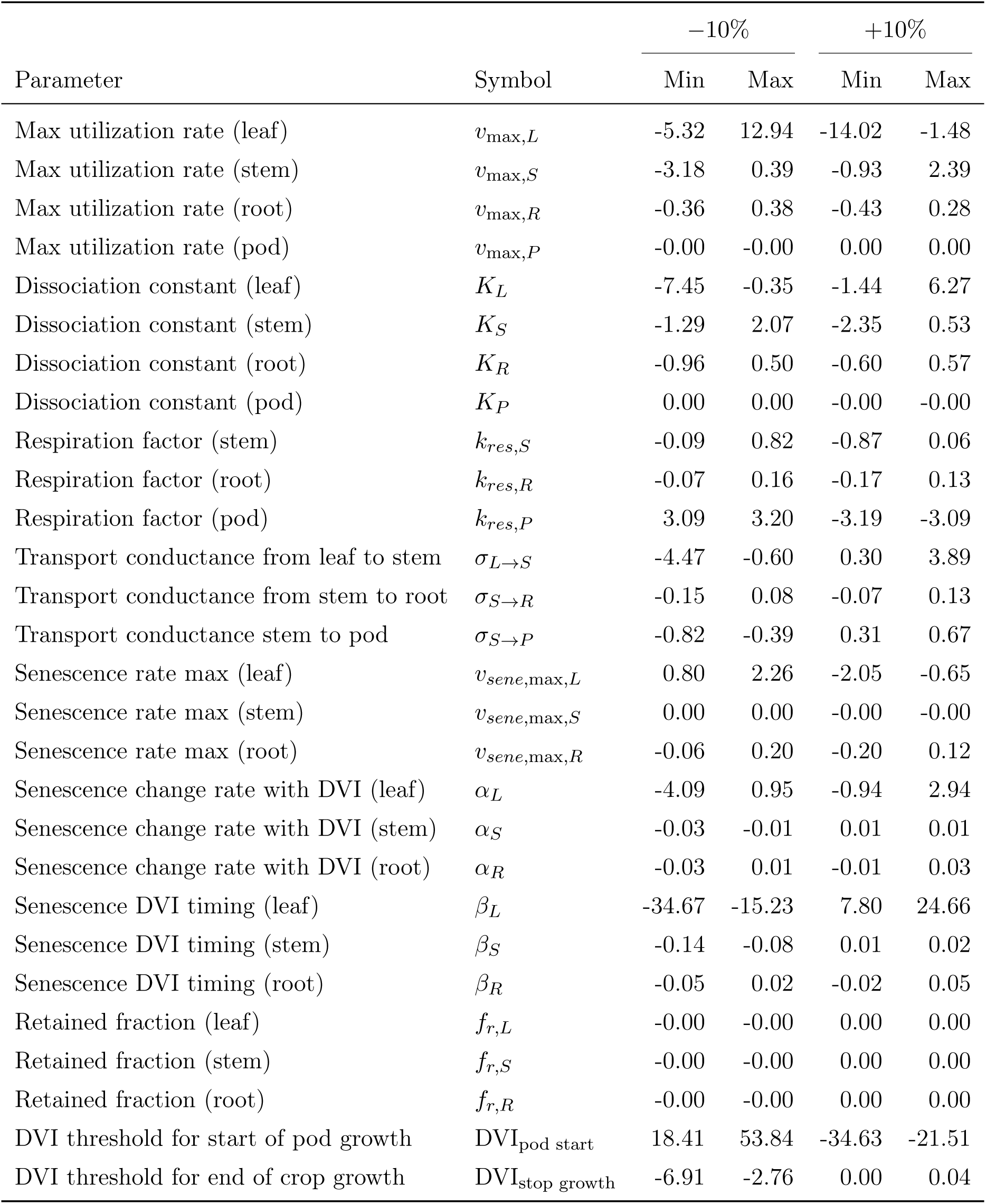
Percent change in final pod mass from a *±*10% UTR parameter perturbation (min – max across 12 cultivar *×* year *×* CO_2_ combinations). All percentages are rounded to two decimal places.

| Parameter | Symbol | –10% |  | +10% |  |
| --- | --- | --- | --- | --- | --- |
|  |  | Min | Max | Min | Max |
| Max utilization rate (leaf) | $v_{\max,L}$ | -5.32 | 12.94 | -14.02 | -1.48 |
| Max utilization rate (stem) | $v_{\max,S}$ | -3.18 | 0.39 | -0.93 | 2.39 |
| Max utilization rate (root) | $v_{\max,R}$ | -0.36 | 0.38 | -0.43 | 0.28 |
| Max utilization rate (pod) | $v_{\max,P}$ | -0.00 | -0.00 | 0.00 | 0.00 |
| Dissociation constant (leaf) | $K_L$ | -7.45 | -0.35 | -1.44 | 6.27 |
| Dissociation constant (stem) | $K_S$ | -1.29 | 2.07 | -2.35 | 0.53 |
| Dissociation constant (root) | $K_R$ | -0.96 | 0.50 | -0.60 | 0.57 |
| Dissociation constant (pod) | $K_P$ | 0.00 | 0.00 | -0.00 | -0.00 |
| Respiration factor (stem) | $k_{res,S}$ | -0.09 | 0.82 | -0.87 | 0.06 |
| Respiration factor (root) | $k_{res,R}$ | -0.07 | 0.16 | -0.17 | 0.13 |
| Respiration factor (pod) | $k_{res,P}$ | 3.09 | 3.20 | -3.19 | -3.09 |
| Transport conductance from leaf to stem | $\sigma_{L \rightarrow S}$ | -4.47 | -0.60 | 0.30 | 3.89 |
| Transport conductance from stem to root | $\sigma_{S \rightarrow R}$ | -0.15 | 0.08 | -0.07 | 0.13 |
| Transport conductance stem to pod | $\sigma_{S \rightarrow P}$ | -0.82 | -0.39 | 0.31 | 0.67 |
| Senescence rate max (leaf) | $v_{sene,\max,L}$ | 0.80 | 2.26 | -2.05 | -0.65 |
| Senescence rate max (stem) | $v_{sene,\max,S}$ | 0.00 | 0.00 | -0.00 | -0.00 |
| Senescence rate max (root) | $v_{sene,\max,R}$ | -0.06 | 0.20 | -0.20 | 0.12 |
| Senescence change rate with DVI (leaf) | $\alpha_L$ | -4.09 | 0.95 | -0.94 | 2.94 |
| Senescence change rate with DVI (stem) | $\alpha_S$ | -0.03 | -0.01 | 0.01 | 0.01 |
| Senescence change rate with DVI (root) | $\alpha_R$ | -0.03 | 0.01 | -0.01 | 0.03 |
| Senescence DVI timing (leaf) | $\beta_L$ | -34.67 | -15.23 | 7.80 | 24.66 |
| Senescence DVI timing (stem) | $\beta_S$ | -0.14 | -0.08 | 0.01 | 0.02 |
| Senescence DVI timing (root) | $\beta_R$ | -0.05 | 0.02 | -0.02 | 0.05 |
| Retained fraction (leaf) | $f_{r,L}$ | -0.00 | -0.00 | 0.00 | 0.00 |
| Retained fraction (stem) | $f_{r,S}$ | -0.00 | -0.00 | 0.00 | 0.00 |
| Retained fraction (root) | $f_{r,R}$ | -0.00 | -0.00 | 0.00 | 0.00 |
| DVI threshold for start of pod growth | DVI<sub>pod start</sub> | 18.41 | 53.84 | -34.63 | -21.51 |
| DVI threshold for end of crop growth | DVI<sub>stop growth</sub> | -6.91 | -2.76 | 0.00 | 0.04 |

The yield was not as sensitive to pod utilization parameters compared to those of the vegetative organs. Final pod mass was most sensitive to the leaf utilization parameters. Increasing the leaf utilization rate decreased the final pod mass in all scenarios (−1.5% to −14.0%). Reducing vegetative organ utilization rates, achieved either by decreasing the maximum utilization rate (*r*_max_) or by increasing the dissociation constant (*K*_O_), had mixed effects. With a 10% lower leaf maximum utilization rate, the final pod mass changed by −5.3% to +12.9%. With a 10% higher leaf dissociation constant, the final pod mass changed by −1.4% to +6.3%. Changes in stem and root utilization, respiration, and transport parameters produced smaller mixed effects (Table 1).

### 3.4 The UTR-BioCro model predicts carbohydrate concentration dynamics in soybean leaves and stems

The LD11-2170 substrate C, measured as total non-structural carbohydrates (TNC) and expressed as moles C per dry mass, averaged 3.5 *±* 2.5 mol kg^-1^ (10.6 *±* 7.5 % dry mass) in the leaf and 2.7 *±* 1.6 mol kg^-1^ (8.0 *±* 4.7% dry mass) in the stem, averaged across all timepoints (Figures 6-7, Tables S4–S5), consistent with previously reported values (Ainsworth et al., 2006; Allen et al., 1998; Ciha & Brun, 1978; Hussain et al., 2019; Liu et al., 2004; Rogers et al., 2004; Streeter & Jeffers, 1979). Inter-plant variability was high at each sampling time, with coefficients of variation (CV) reaching up to 47% for leaves and 71% for stems (Tables S4-S5), reflecting high variability in plant substrate carbon concentrations. The model captures the higher concentration in leaves than stems at most sampling times, and the diurnal pattern of daytime accumulation and nighttime drawdown, including the larger diurnal amplitude in leaves than stems (Figure 7).

**Figure 6:**
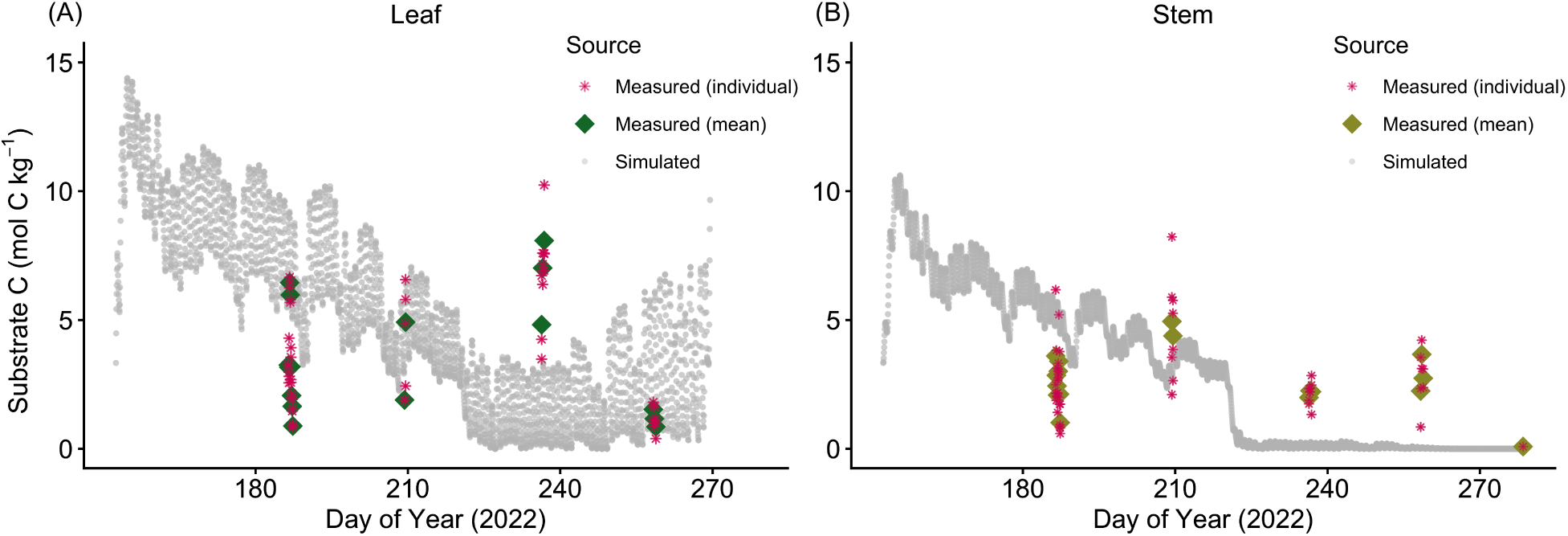
Measured and predicted substrate C concentrations (mol C kg*^−^*^1^ dry mass) in LD11-2170 (A) leaves and (B) stems throughout the 2022 growing season. Substrate C was estimated as TNC (total nonstructural carbohydrates) measured from the top fully expanded leaf and its closest stem section. At each time point, four plants were sampled; technical replicates with a coefficient of variation exceeding 10% were excluded.

**Figure 7:**
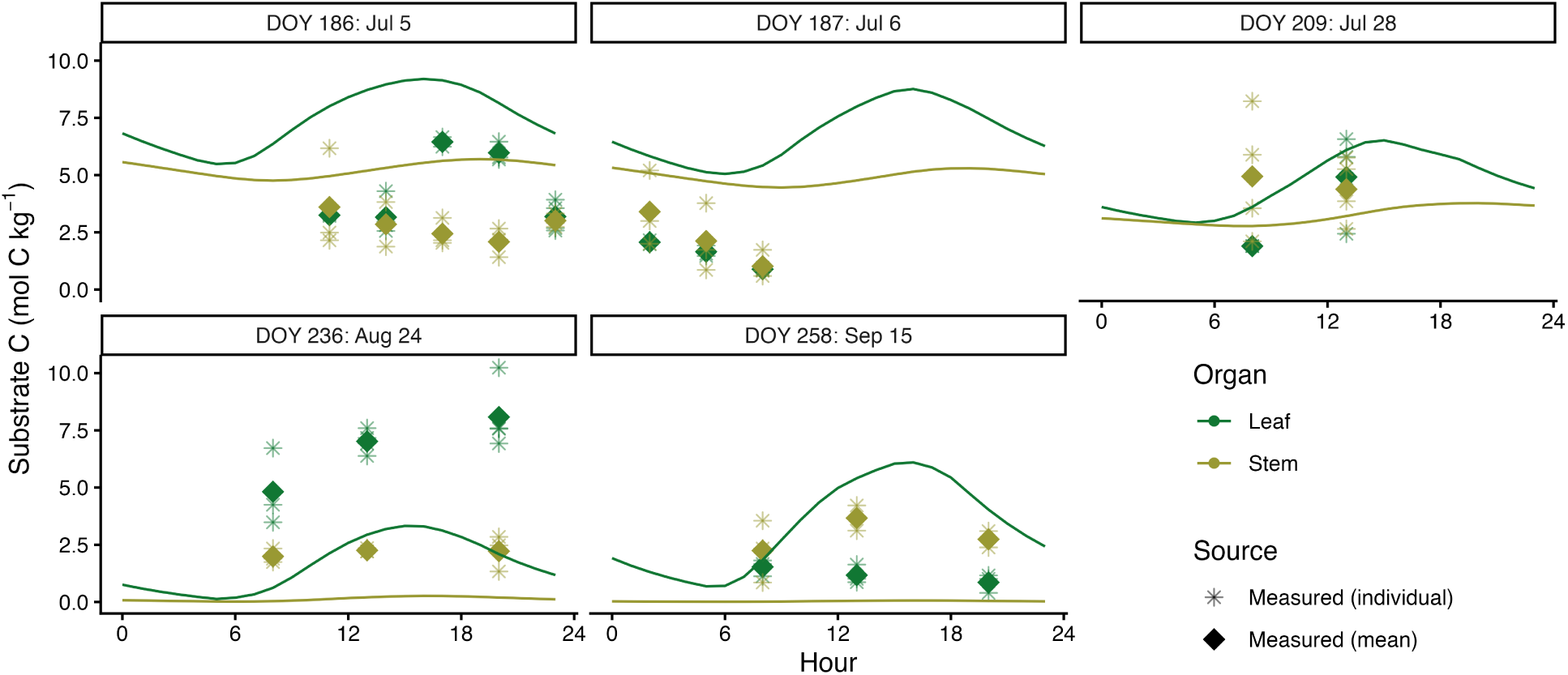
Diurnal dynamics of substrate C concentration (mol C kg*^−^*^1^ dry mass) in LD11-2170 leaves and stems on the sampling days in 2022, compared with UTR-BioCro predictions. At each time point, four plants were sampled; technical replicates with a coefficient of variation exceeding 10% were excluded.

Model predictions differed from observations on August 24 (DOY 236) and September 15 (DOY 258). The measured TNC on August 24 (growth stages V16 and R6) for both leaf and stem exceeded model predictions. This difference could be due to the differences between the TNC at the top of the canopy where the measurements were taken compared to the average canopy TNC which is what the model predicts. Compared to the earlier measurements, the canopy closure at this stage causes substantial self-shading, creating large gradients in photosynthetic rates between the top (7.0 µmol m^-2^ s^-1^) and bottom (1.0 µmol m^-2^ s^-1^) of the canopy (Figure S3). More photosynthetically active leaves at the top of the canopy would accumulate more TNC, producing a vertical TNC gradient that the current UTR model does not capture. Additionally, the leaf TNC consisted of a larger percentage of starch (89–94%) on August 24 than the average (56–88%) measured on the other sampling days (Table S4). TNC is used as a proxy for substrate C, but the two are not equivalent: soluble sugar, rather than starch, is the more mobile and readily available carbon pool, and so more closely matches the model’s definition of substrate C. Because the UTR model does not represent starch explicitly, it cannot capture this starch buildup, which may explain why the model’s predicted substrate C falls below the measured TNC concentration on Aug 24.

On September 15, the observed leaf TNC was lower than stem TNC, contrary to the model prediction. We attribute this discrepancy to a decline in photosynthetic capacity accompanying leaf senescence, a process not currently represented in the UTR-BioCro model. Visibly brown leaf samples collected on this date indicate chlorophyll degradation and the associated reduction in assimilation rate, which is consistent with the observed drop in leaf TNC.

### 3.5 UTR-BioCro model predicts yield responses to source-sink manipulation

We assessed the ability of the UTR-BioCro model to capture changes in how carbon is allocated by simulating crop responses to different source-sink perturbations. The manipulations include altering source strength (canopy shading), sink strength (pod removal), and both source and sink strengths (defoliation).

#### 3.5.1 Shading

Across the four shading levels, the experimentally observed yield reduction rose gradually from 18.1% at 50% shade to 37.9% at 80% shade, increasing more slowly than the imposed shading percentage (Proulx & Naeve, 2009). The UTR-BioCro model tracked this gradual response, predicting reductions of 18.5% at 50% to 32.3% at 80% shade respectively (Figure 8). In contrast, the partitioning-BioCro model overpredicted yield reduction at every shading level (33.7% to 60.5%), and the gap between the two models widened as shading intensified. In the partitioning model, yield reduction closely mirrored the shading percentage, as nearly all net assimilation is directed to seed by R5.

**Figure 8:**
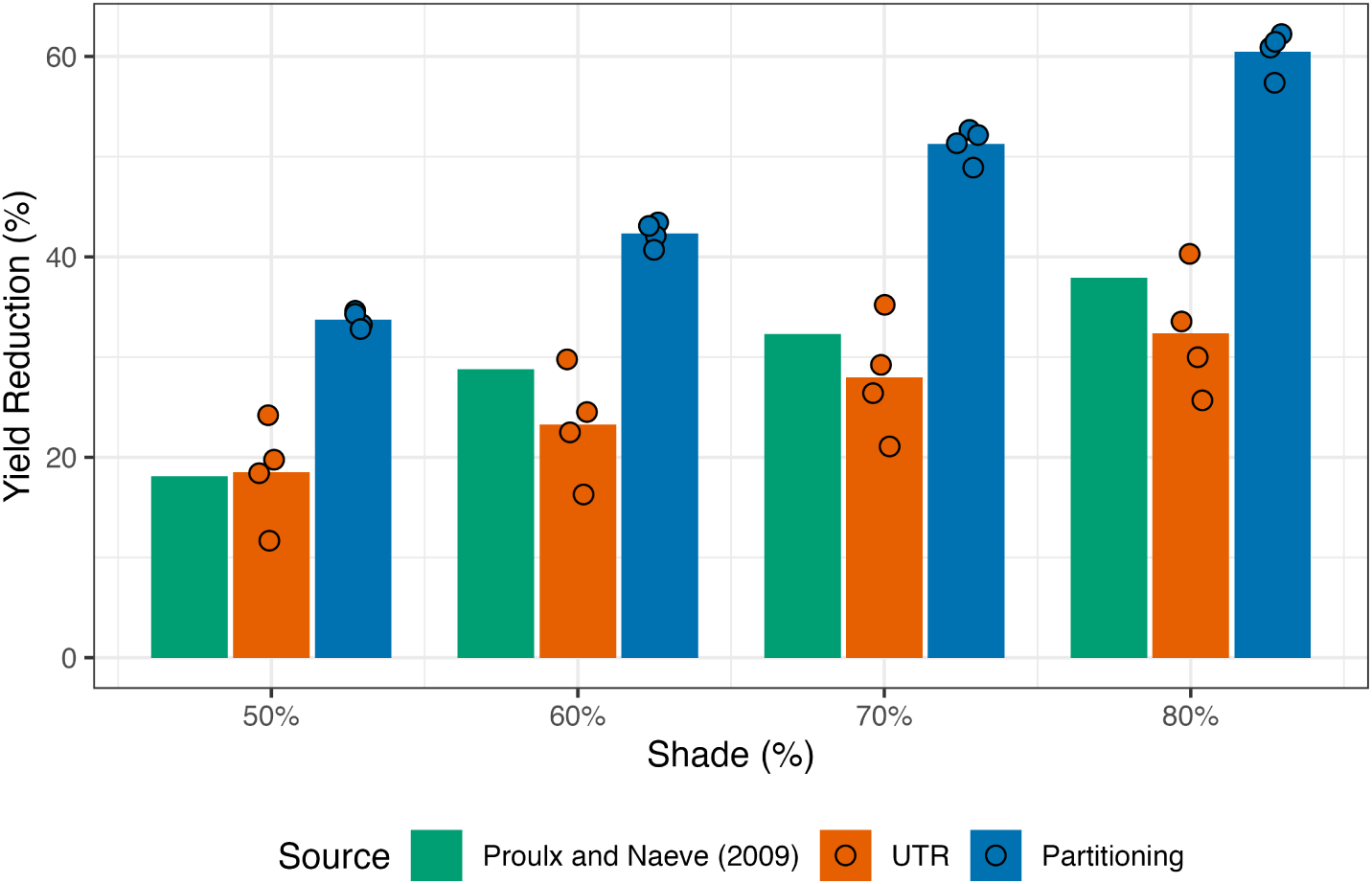
Predicted and observed (Proulx & Naeve, 2009) reductions in yield under 50%, 60%, 70%, and 80% shading applied from R5 (DVI=1.5) to the end of the growing season. Bars represent average reduction percentages; dots represent individual year simulations (2002, 2004–2006) at ambient CO_2_.

#### 3.5.2 Pod removal

Observed yield reductions increased from 11.7% at 20% pod removal to 52.3% at 70% removal (Proulx & Naeve, 2009). The UTR-BioCro model predicted similar reductions in yield (15.9% to 55.7%) across all four removal levels, within 5% away from the experimental means (Figure 9). The partitioning-BioCro model only predicted a similar decrease in yield (10.5%) as the observed for the 20% pod removal level. At the higher pod removal levels, the partitioning-BioCro model consistently underpredicted the yield response, diverging further from the observations as more pods were removed.

**Figure 9:**
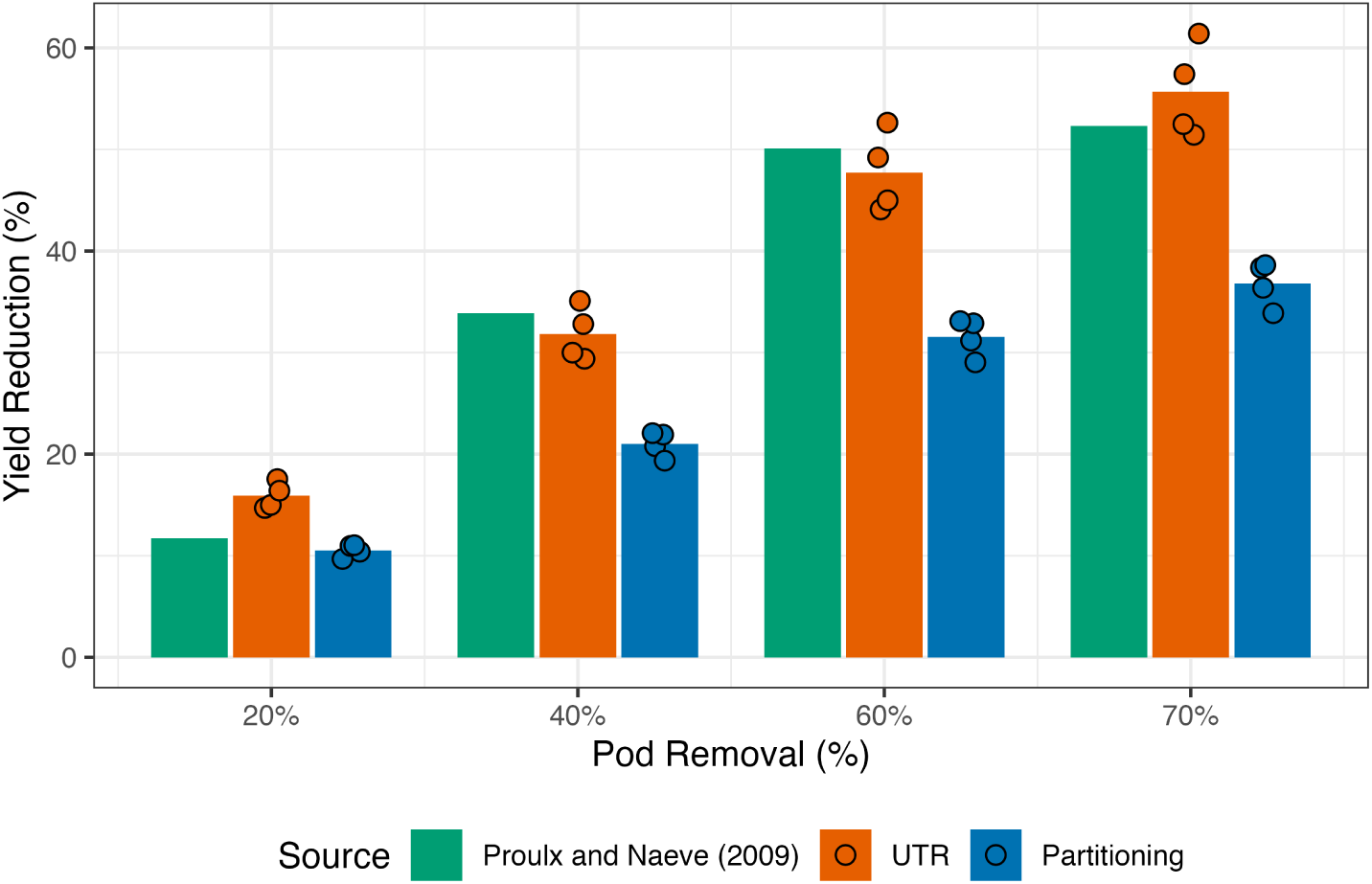
Predicted and observed (Proulx & Naeve, 2009) reductions in yield from 20%, 40%, 60%, and 70% pod removal at R5 (DVI=1.5). Bars represent average reduction percentages; dots indicate individual year simulations (2002, 2004–2006) at ambient CO_2_.

#### 3.5.3 Defoliation

##### Defoliation during reproductive stages

In the defoliation experiments of Parvej et al. (2025), yield reduction increased from near zero at no defoliation to 73.2% (Stage R4) and 81.7% (Stage R5) at full defoliation. At R4, the UTR-BioCro model predicted reductions in yield of 0.6% to 65.3% compared to observed values of 7.0% to 73.2% at 25% to full defoliation (Figure 10A). The UTR-BioCro model tracked the increase in yield reduction across defoliation levels but underpredicted at intermediate defoliation intensities. The partitioning-BioCro model also underpredicted yield reduction from 0–75% defoliation, even predicting a small yield gain at 25% defoliation (−1.3% reduction), but overpredicted at full defoliation (83.2% vs. 73.2% observed yield reduction). At R5, both models underpredicted the yield reductions across all defoliation levels (Figure 10B). The UTR-BioCro model predicted only a 36.3% yield reduction against an observed 81.7% at full defoliation. The partitioning-BioCro model was closer to the experimental result at full defoliation at R5 (64.0% versus 81.7%), but the two models both underpredicted the impact on yield at lower defoliation levels.

**Figure 10:**
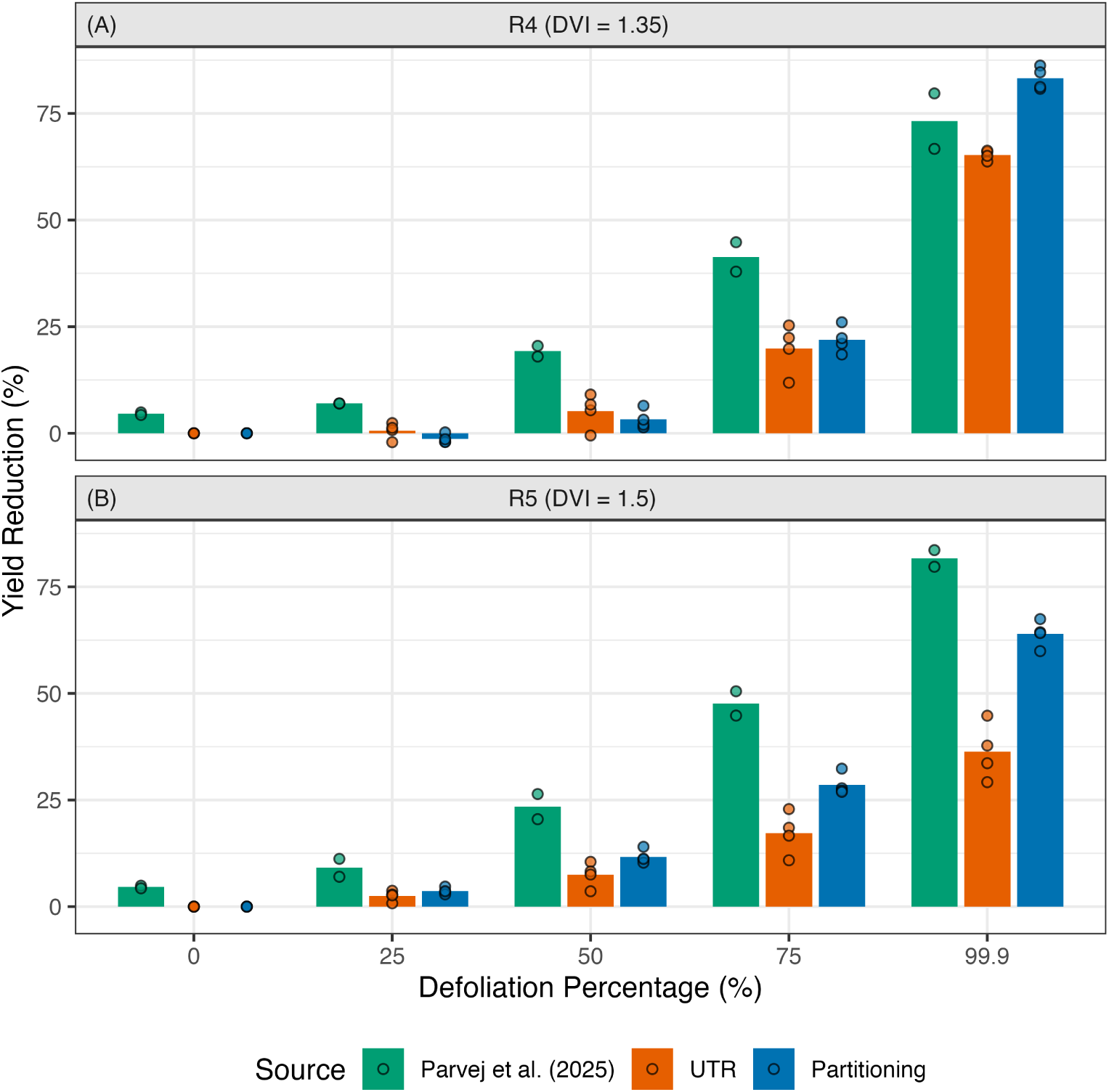
Predicted and observed (Parvej et al., 2025) reductions in yield from 0, 25%, 50%, 75%, and 100% (99.9% in the model) defoliation at (A) Stage R4 (DVI=1.35) and (B) Stage R5 (DVI=1.5). Bars represent average yield reduction percentages. Dots for the experimental data show the yield reductions in Iowa and Indiana. Dots for the model results show the yield reductions from individual years (2002, 2004–2006) at ambient CO_2_.

##### Defoliation due to hail event

A hail event occurred on DOY 198 in 2003 during the vegetative stage (V7) of the SoyFACE experiments that caused 60% leaf loss and 21% aboveground biomass loss (Morgan et al., 2005). Neither model handles this unusual event well, and this is an extreme test of a model’s predictive ability. However, compared to the partitioning model, the UTR-BioCro model more closely reproduces leaf and stem biomass after the hail event under both ambient and elevated CO_2_, without requiring parameter adjustments (Figure 11). Following the simulated loss, predicted leaf and stem biomass remained suppressed and recovered slowly, matching the trajectory of the measured data. The partitioning-BioCro model instead re-grew leaf and stem mass rapidly, overpredicting both organs (Figure S5). The UTR-BioCro model captured the slower post-hail growth because the utilization rate declined with structural mass (Figure S6), while re-growth in the partitioning model is substantially faster (Figure S5). However, both models overpredict the final pod mass. The UTR-BioCro model overpredicts final pod biomass by approximately 40% under both ambient and elevated CO_2_ in the 2003 hail simulation, likely because the model does not account for delayed onset of pod filling (Kalton et al., 1949).

**Figure 11:**
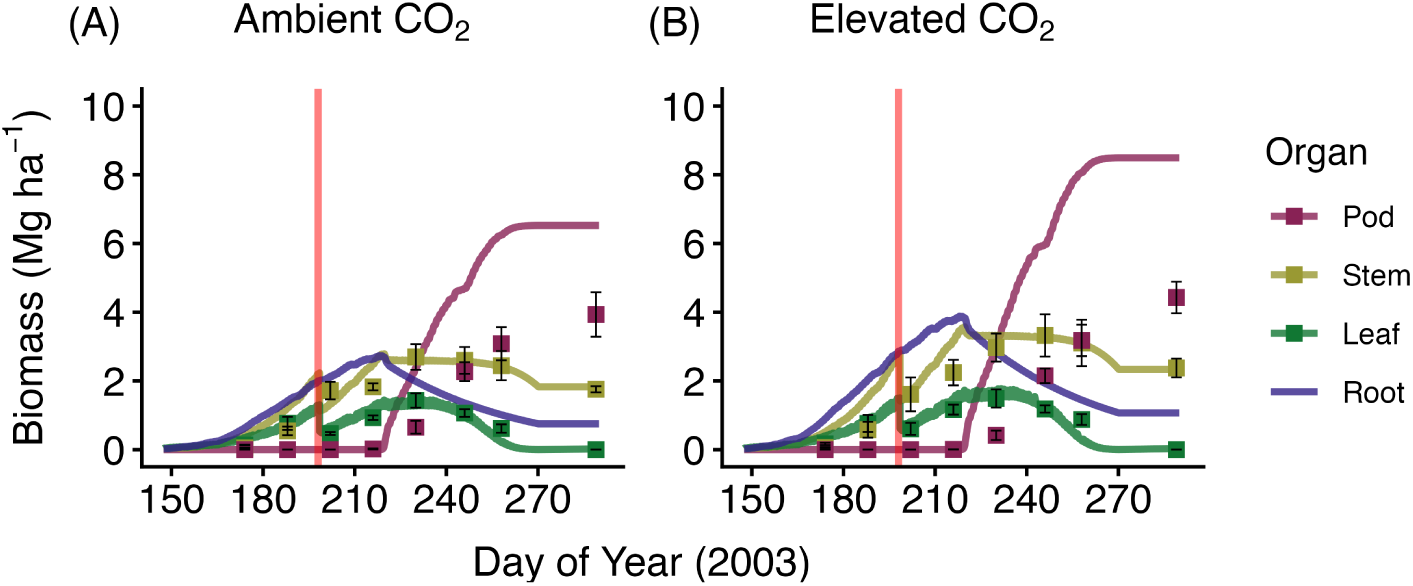
Simulated soybean growth compared to biomass data in 2003 with a hail event on DOY 198 (denoted by the vertical red bar) at (A) ambient and (B) elevated CO_2_ levels using the UTR model.

## 4 Discussion

### 4.1 Emergent source-sink responses of the UTR framework

A key advantage of the process-based UTR framework is that crop responses to source–sink perturbations emerge from local utilization and transport dynamics rather than from prescribed allocation rules. None of the elevated CO_2_, shading, pod removal, defoliation, or hail scenarios were used during calibration; the model parameters were fit using only soybean organ biomasses at ambient CO_2_ in 2002 and 2005. Despite this, the UTR-BioCro model reproduces yield reduction under shading and pod removal more closely than the partitioning model (Figures 8, 9), and reproduces post-hail leaf and stem trajectories that the partitioning model overpredicts (Figure 11, Figure S5).

When simulating shading, the UTR model predicted a smaller yield reduction than the partitioning model, accurately capturing the measured reductions in field experiments (Proulx & Naeve, 2009). The UTR model predicts a smaller reduction in yield because the sink strength of the pod is not changed by the shading, creating a similar substrate C gradient which drives the transport of C from leaf to stem and to pod, even though the source strength is reduced. Similarly, when simulating pod removal, the UTR model accurately predicted a higher yield reduction compared to the partitioning model because the UTR model captures the resulting reductions in carbon utilization and transport to the pod, which the partitioning approach ignores.

These results demonstrate that a simple set of process-based equations, Hill-equation based utilization rates, gradient-driven transport rates, and DVI-controlled organ activation, can reproduce the qualitative and often quantitative response of soybean to a wide range of source-sink perturbations without any scenario-specific tuning. This is a substantive step beyond partitioning-based crop models, including the teleonomic and sink competition models, which require new parameter sets or explicit stress modules to handle each scenario.

### 4.2 Current limitations in modeling defoliation treatments

The UTR model’s underestimation of defoliation-induced yield loss (Figures 10&11) likely reflects several processes the model omits: pod abortion (Board et al., 2010; Board & Tan, 1995; Carrera et al., 2022; Fehr et al., 1977; Poudel et al., 2025), reduced seed size (Board et al., 2010; Carrera et al., 2022; Poudel et al., 2025), and a shortened seed-filling period (Fehr et al., 1977). In particular, a shortened seed-filling period and reduced seed size would likely decrease the seed:pod ratio which we assume to be invariant between treatments in our analysis. If we were to assume a smaller seed:pod ratio for the defoliation treatments, then the predicted yield reductions from the UTR-BioCro model would be larger, more closely reflecting what was observed. Similarly, pod abortion would decrease the sink strength of the pod which would also result in larger predicted yield reductions. Separating the seed and shell into different organs and incorporating a pod abortion module could help address some of these limitations. Further, better mechanistic understanding and incorporation of how crop development is impacted by defoliation events would further address limitations and improve yield predictions in response to damaging incidents such as the hail event in 2003 (Figure 11).

### 4.3 A mechanistic hypothesis for the increase in root:shoot ratio under elevated CO_2_

The UTR model offers a mechanistic hypothesis for the increase in R/S under elevated CO_2_, a phenomenon traditionally explained by the functional balance principle, where carbon allocation aims to balance capacity to obtain different resources (Le Roux et al., 2001) (Figure 5). In the UTR framework, elevated CO_2_ increases substrate carbon concentrations, driving utilization rates closer to saturation (Figure S8). The “saturation” hypothesis is supported by the observation that concentrations of central metabolic intermediates mostly exceeds their corresponding *K_m_* values, indicating that these intermediates operate in a near-saturating regime (Bennett et al., 2009). Meanwhile, substrate C transport increases linearly with the source-sink concentration gradient. The combined effect is a greater proportion of assimilates being transported to roots (Figure S9), increasing R/S. Notably, this response emerges from carbon dynamics alone, without invoking nutrient feedback or photosynthetic down-regulation attributed to carbohydrate accumulation (Drake et al., 1997).

However, experimental evidence on whether elevated CO_2_ increases soybean R/S is mixed. Some studies report an increased R/S under elevated CO_2_ (Rogers et al., 1992; Rogers et al., 1995), while others find no significant change (Ainsworth et al., 2002; Idso et al., 1988). This inconsistency suggests that additional processes may counteract or obscure the carbon-driven increase in R/S under elevated CO_2_. Three potential mechanisms have been suggested to explain why R/S doesn’t always increase under elevated CO_2_. First, elevated CO_2_ reduces stomatal conductance, lowering canopy transpiration (Ainsworth et al., 2002), thereby alleviating water stress and weakening the stimulus for root growth, which dampens the R/S response. Second, the higher sucrose concentrations at elevated CO_2_ can down-regulate the expression of sucrose transporters (Chiou & Bush, 1998). Reduced surcrose transport would limit the additional carbon reaching the roots, dampening the R/S response that the UTR model predicts from the concentration gradient alone. Third, elevated sucrose concentrations also increase phloem sap viscosity, which raises the resistance of long-distance transport (Stanfield & Bartlett, 2022). Greater transport resistance reduces carbon flux to roots, again diminishing the increase in R/S. The UTR model represents only the carbon-concentration-gradient contribution to transport and omits these regulatory and physical feedbacks. By isolating the contribution of TNC dynamics, the current UTR framework provides a baseline against which these additional mechanisms could be evaluated.

### 4.4 The UTR model predicts biologically meaningful carbon allocation patterns as emergent properties

The UTR model yields more biologically meaningful carbon allocation fractions, whereas the prescribed partitioning approaches are often constrained by their choice of mathematical forms. The sigmoidal partitioning functions, for example, force root allocation to zero after the mid-vegetative stages (DVI>0.5) and leaf and stem allocation to zero once pod growth starts (Figure 1C in Matthews et al. (2022)). This is inconsistent with the indeterminate growth of soybean. Development stage surveys from 2021 to 2024 confirmed that LD11-2170 continued vegetative growth, expanding from 7–8 to 15-16 trifoliate leaves during the reproductive phase (Table S6). The UTR model captures this behavior; although leaf and stem allocation fractions decline during pod growth (Figure 3), both organs maintain positive growth rates (Figure S7).

The allocation fractions derived from the UTR model (Figure 3A) are also comparable to experimental measurements. The UTR model predicted 40–50% of the assimilated carbon to be allocated to the root prior to pod growth, consistent with the 43% (derived from Table 3 in Harris et al. (1985)) and 51% (Pate & Herridge, 1978) below-ground allocation estimates obtained from ^14^C labeling studies. Similarly, the predicted leaf carbon allocation fraction from the UTR model averaging approximately 10% (Figure 3B) is consistent with prior literature that found that 10% of the photosynthate accumulated in soybean leaves during the vegetative phases with the remaining 90% being lost through respiration and export to other organs (Figure 5 in Rogers et al. (2004)). While both models reproduce organ dry mass growth trajectories, the UTR model produces allocation patterns more consistent with observed soybean physiology.

### 4.5 Developmental transition timing emerges as a potential target for crop improvement

The sensitivity analysis of the UTR model (Table 1) reveals that final pod mass is most sensitive to parameters governing developmental stage transitions. This underscores the importance of accurately modeling transition timing, which depends on both environmental conditions (Figure 11) (Desclaux & Roumet, 1996) and the cultivar. The result is consistent with the observation that the duration of seed filling is positively correlated with seed yield (Dunphy et al., 1979; Hanway & Weber, 1971; Nelson, 1986) and has been used successfully as a screening criterion for yield improvement (Zeinali-khanghah et al., 1993). In contrast, perturbations to utilization and transport parameters produce mixed effects on final pod mass (Table 1). This may explain why attempts to increase yield by overexpressing sucrose transporters, at either the source or the sink, have generally not produced significant gains (Ainsworth & Bush, 2011; Leggewie et al., 2003; Weichert et al., 2010). The contrasting sensitivity suggests that transition timing may be a more effective target for yield improvement.

### 4.6 Organ-level TNC representation enables future coupling to metabolic models and signaling pathways

A mechanistic description of organ-level carbon dynamics is also advantageous for using crop models to evaluate engineering and breeding strategies. Photosynthesis is typically the only metabolic process mechanistically described in crop models, while other processes, including carbon allocation, remain empirically represented, limiting models’ ability to project how organ-specific metabolic modifications propagate to whole-plant growth and yield (Piao & Matthews, 2026). Because the UTR model’s utilization and respiration parameters reflect organ-specific metabolic costs, they can be modified directly to explore the potential systemic consequences of an engineering intervention, rather than requiring re-calibration of partitioning fractions whose connection to the underlying biology is indirect. The explicit organ-level representation of substrate C opens up several such modeling possibilities that extend beyond the scope of the current work. For example, the model could be used to evaluate the tradeoff between soybean yield and oil content by changing the pod utilization and respiration parameters based on the different C costs to produce protein, oil, and residuals (Hanson et al., 1961). The framework could also be applied to study the impacts of modified traits in other crops, such as enhanced lipid production in leaves and stems of bioenergy crops (Chen et al., 2026; Clark & Schwender, 2022), which would raise utilization and respiration rates.

Since the UTR-BioCro framework resolves substrate C concentration in each organ at an hourly resolution, these concentrations could also be used as inputs to sugar-signaling responses within the plant. For example, high leaf TNC down-regulates photosynthesis through feedback inhibition (Drake et al., 1997), elevated hexose concentrations promote leaf senescence (Wingler et al., 2006), while sugar starvation triggers both senescence (Wingler et al., 2006) and reproductive abscission (Antos & Wiebold, 1984). Similarly, accelerated leaf senescence under prolonged source limitation, or pod abortion under source–sink imbalance (Board & Tan, 1995; Egli & Bruening, 2005) could be incorporated through describing senescence parameters as functions of substrate C concentrations. Currently, these processes depend only on the environment and predicted developmental stage, without endogenous input from sugar-signaling.

Additionally, the UTR-BioCro framework could also represent organ-specific environmental responses since each organ has its own substrate pool and utilization parameters. For example, maize ear components differ in their heat sensitivity (Suwa et al., 2010), and leaf growth is inhibited more strongly than root growth under water stress (Hsiao & Xu, 2000). In the UTR formulation, such effects could be included by making the impacted organ’s utilization rate *r*_max,O_ a function of the local stressor, with the consequences for whole-plant growth and allocation emerging from the same set of equations. Because these parameters correspond more directly to specific enzymes and anatomical traits than partitioning coefficients do, the UTR framework enables are more straightforward approach to pose mechanistic questions, such as how increasing oil content would affect whole-plant carbon allocation and yield. Such questions are difficult to be answered in partitioning-based models, where organ-specific responses require coordinated and often ad hoc adjustments to multiple allocation coefficients to preserve mass balance. Together, these capabilities point toward a modeling framework in which whole-plant growth and yield emerge from organ-level carbon dynamics, signaling, and stress physiology.

## 5 Conclusion

In this study, we integrated a UTR carbon allocation model into the Soybean-BioCro crop growth modeling framework. We adapted this UTR framework from Thornley’s original model (Thornley, 1972), to work with the BioCro model and extended the UTR framework by adding reproductive growth and senescence. We validated the model against organ biomass from two cultivars grown across two CO_2_ levels over eight growing seasons, achieving accuracy comparable to partitioning models. Predicted leaf and stem TNC concentrations also aligned reasonably with experimental measurements. A local sensitivity analysis of model parameters identified developmental transition timing, particularly the onset of pod growth and leaf senescence, as having the strongest impact on final yield, while perturbations to utilization and transport parameters had mixed effects. This identifies phenological timing as a promising target for both crop improvement.

A key strength of this UTR-BioCro framework is that biologically meaningful behaviors emerge without scenario-specific tuning. Carbon allocation fractions, vegetative-stage storage and reproductive-stage mobilization, the increase in root:shoot ratio under elevated CO_2_, and yield responses to shading, pod removal, defoliation, and hail damage all arise from the same set of process-based equations. Capturing this range of responses with a single calibration distinguishes the approach from partitioning models, which typically require new parameter sets or dedicated stress modules for each scenario. It also avoids the “optimizing” assumptions underlying teleonomic models and redefining the “growth potentials” for each scenario in sink competition models. The UTR framework also offers a starting point for incorporating nutrient allocation, sugar signaling, and organ-specific stress responses by linking organ-level substrate carbon dynamics to whole-plant growth. This approach could strengthen the predictive capacity of crop models under novel environments and genetic backgrounds, supporting the development of crop improvement strategies.

## Acknowledgments

We thank Scott Oswald for valuable conversations about this work and particularly the source-sink manipulation scenarios.

This work was partially funded by the U.S. Department of Agriculture National Institute of Food and Agriculture under Grant 2020-68012-31674, the Foundation for Food & Agriculture Research under award number Grant ID: 602757, and the DOE Center for Advanced Bioenergy and Bioproducts Innovation (U.S. Department of Energy, Office of Science, Biological and Environmental Research Program under Award Number DE-SC0018420). Any opinions, findings, and conclusions or recommendations expressed in this publication are those of the authors and do not necessarily reflect the views of the US Department of Agriculture (USDA) or other funding agencies. Mention of trade names or commercial products in this publication is solely for the purpose of providing specific information and does not imply recommendation or endorsement by the USDA. USDA is an equal opportunity provider and employer.

## Notes

### Competing Interest Statement

The authors have declared no competing interest.

