## Supplementary Information for "Integrating carbon utilization and transport processes into a crop growth model enables the prediction of emergent soybean carbon allocation behavior"

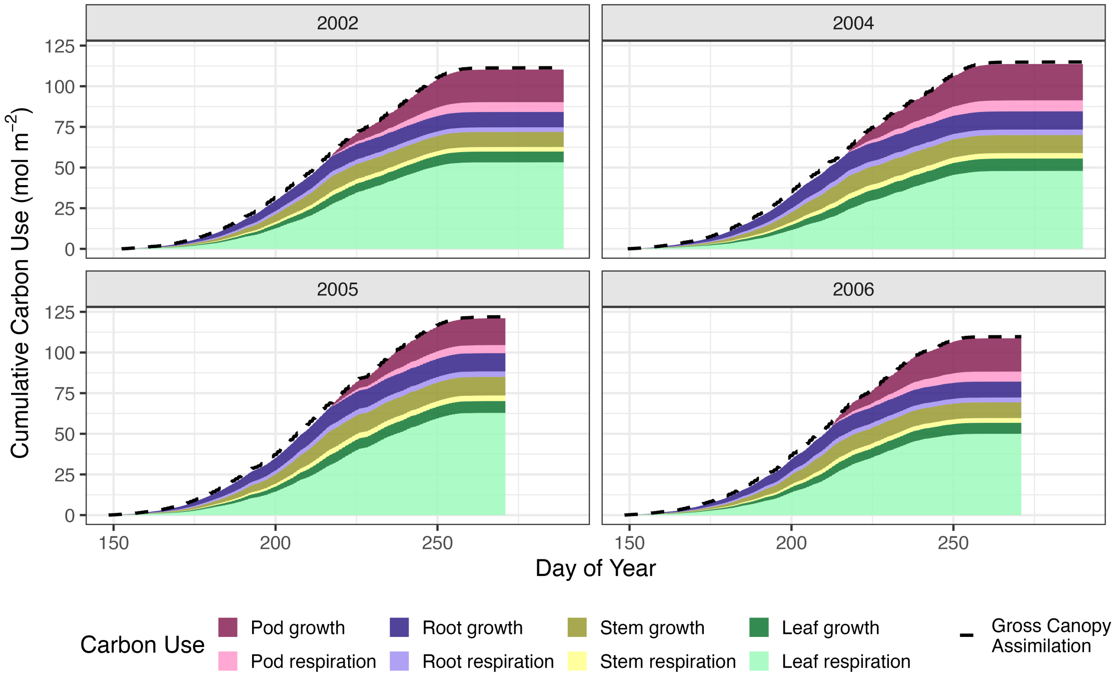


Figure S1: Cumulative carbon use in Pioneer 93B15 at ambient $CO_2$ from the UTR model. The dashed line represents the cumulative net canopy assimilation rate.

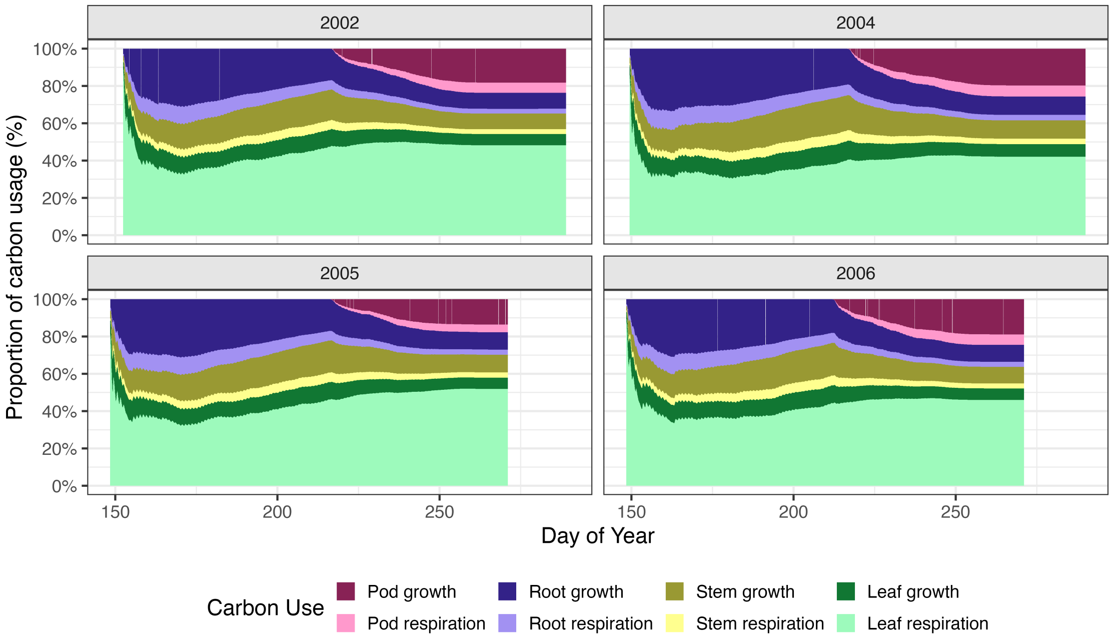


Figure S2: Fractions of organ cumulative carbon use compared to the total use in Pioneer 93B15 at ambient $CO_2$ from the UTR model.

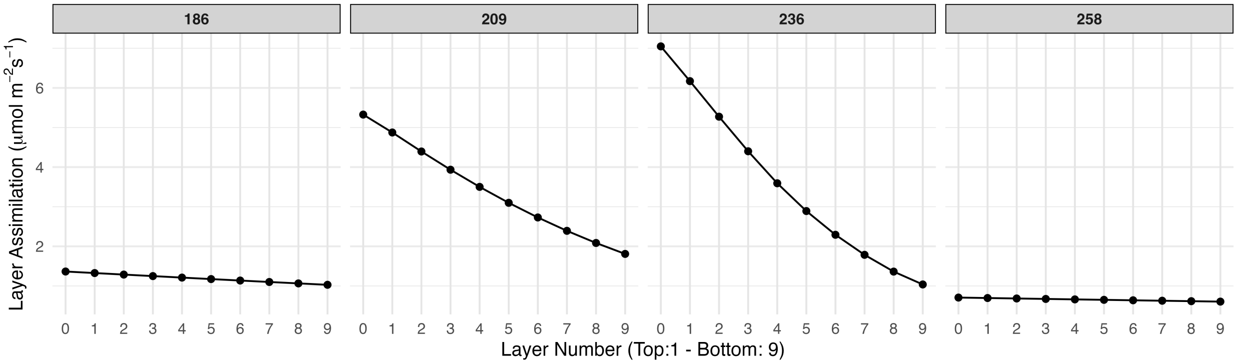


Figure S3: Assimilation rate per ground area (µmol m$^{-2}$ s$^{-1}$) from top to bottom on different DOYs from BioCro.

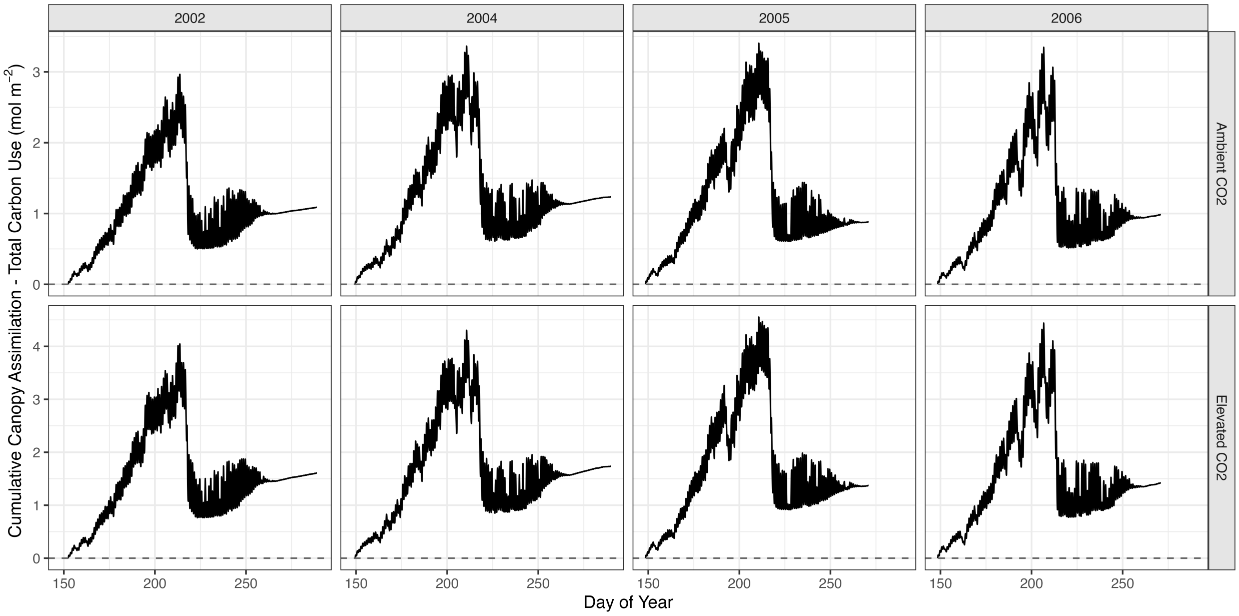


Figure S4: Difference between total canopy assimilation rate and cumulative carbon use in Pioneer 93B15 at ambient and elevated $CO_2$ levels.

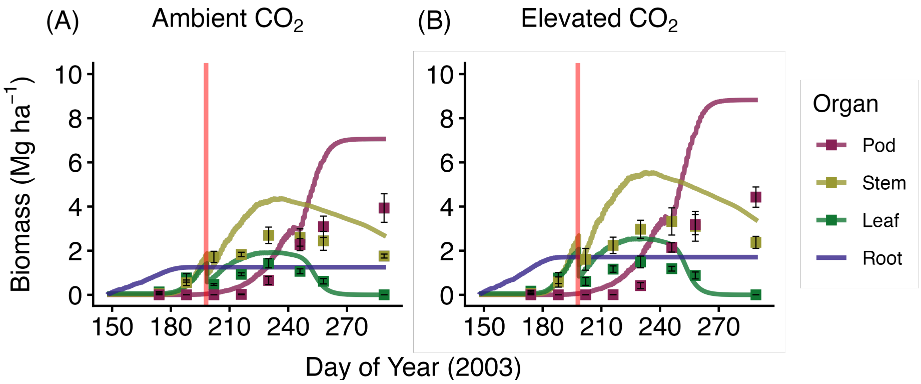


Figure S5: Simulated soybean growth compared to biomass data in 2003 at ambient and elevated $CO_2$ levels using the partitioning model.

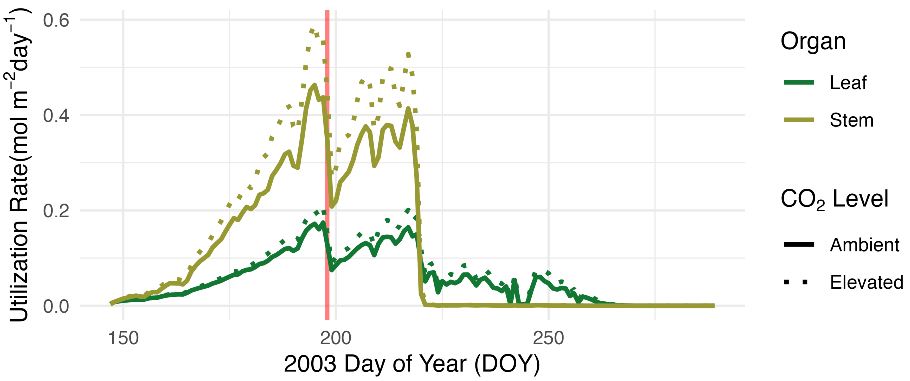


Figure S6: Simulated soybean leaf and stem daily utilization rates in 2003 at ambient and elevated $CO_2$ levels from the UTR-BioCro model. The red line marks the hail event on DOY 198.

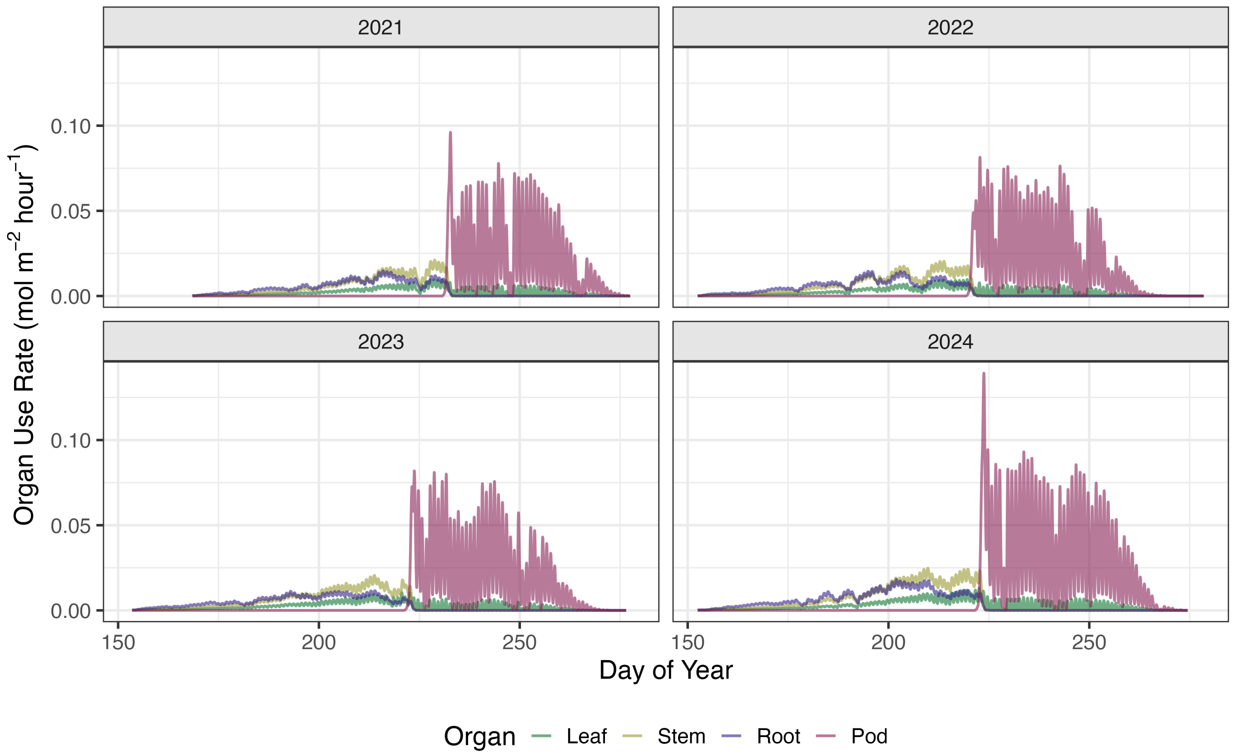


Figure S7: LD11-2170 hourly organ substrate carbon use rates (mol m$^{-2}$ hr$^{-1}$).

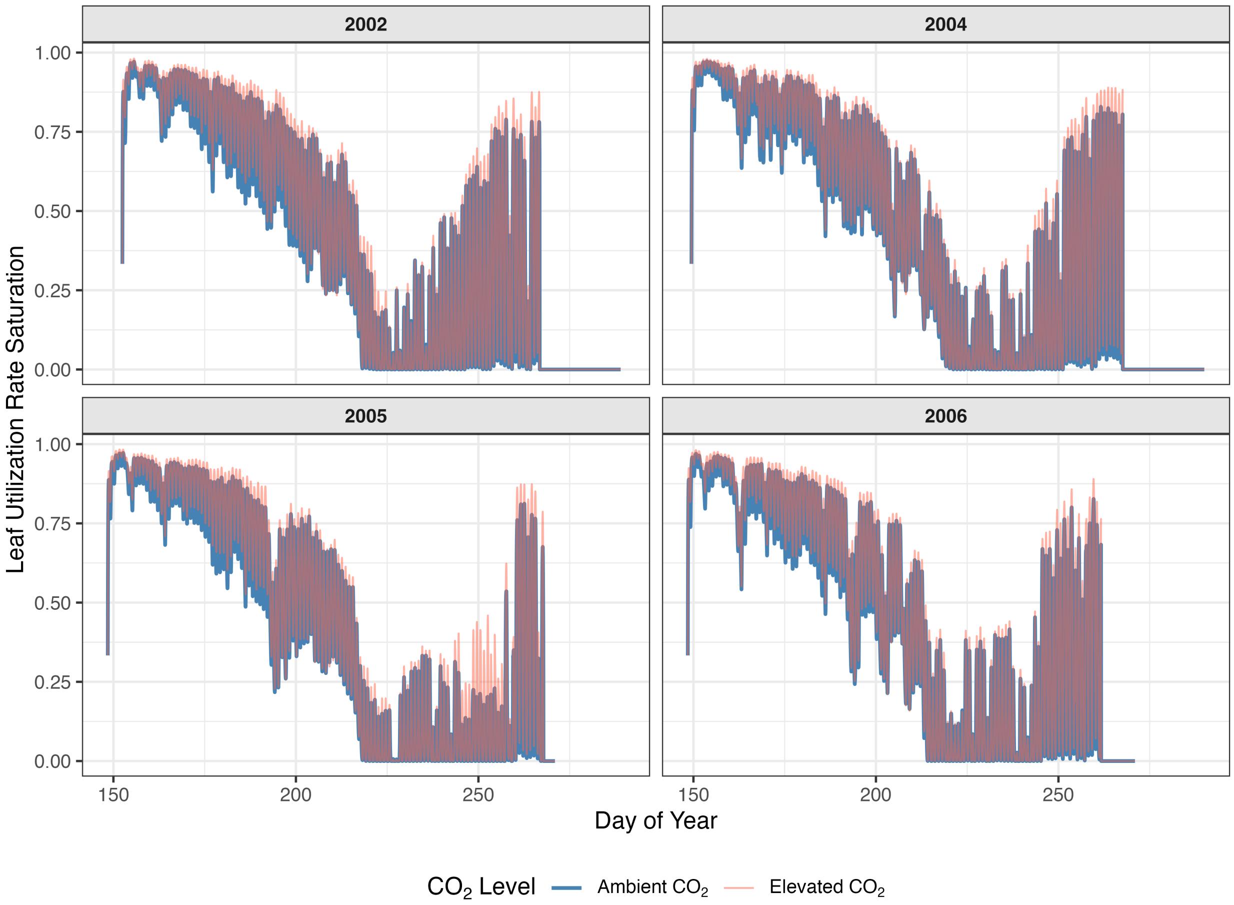


Figure S8: Leaf utilization saturation ratio in ambient $CO_2$ vs elevated $CO_2$ for Pioneer 93B15. Leaf utilization saturation ratio is defined as the ratio of leaf utilization rate per structural C to the maximum utilization rate, $v_{\max,L}$.

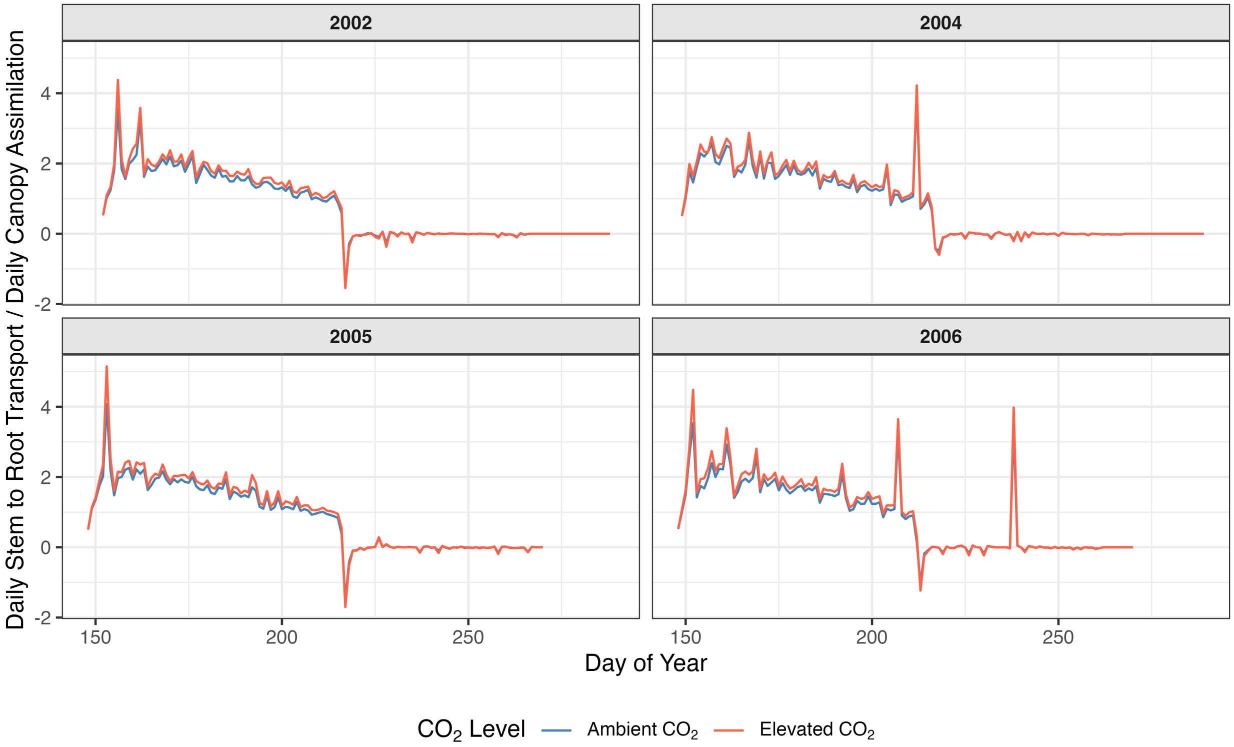


Figure S9: The ratios of daily stem to root substrate transport to daily canopy assimilation rate under ambient $CO_2$ vs elevated $CO_2$ for Pioneer 93B15. The negative values are due to negative net assimilation rate on the corresponding day.

### Supplemental Tables

Table S1: UTR model parameters, estimation ranges, parameterization results, and corresponding equations. L: Leaf. S: Stem. R: Root. P: Pod

| Parameter | Symbol | Unit | Min | Max | Fit | Eq |
|---|---|---|---|---|---|---|
| **Utilization Parameters** | | | | | | |
| Max utilization rate constant (L) | $v_{\max,L}$ | $hr^{-1}$ | 0 | 0.1 | 0.003548 | (3) |
| Max utilization rate constant (S) | $v_{\max,S}$ | $hr^{-1}$ | 0 | 0.1 | 0.009276 | (3) |
| Max utilization rate constant (R) | $v_{\max,R}$ | $hr^{-1}$ | 0 | 0.1 | 0.044412 | (3) |
| Max utilization rate constant (P) | $v_{\max,P}$ | $hr^{-1}$ | 0 | 1 | 0.974662 | (3) |
| Half-saturation constant (L) | $K_L$ | – | 0 | 0.5 | 0.157163 | (3) |
| Half-saturation constant (S) | $K_S$ | – | 0 | 0.5 | 0.205616 | (3) |
| Half-saturation constant (R) | $K_R$ | – | 0 | 0.5 | 0.487155 | (3) |
| Half-saturation constant (P) | $K_P$ | – | 0 | 0.5 | 0.018651 | (3) |
| Respiration fraction (SRP) | $k_{res,O}$ | – | 0.2 | 0.8 | 0.230973 | (4) |
| **Transport Parameters** | | | | | | |
| Transport conductance from leaf to stem | $\sigma_{L\to S}$ | $hr^{-1}$ | 0 | 1 | 0.153715 | (6) |
| Transport conductance from stem to root | $\sigma_{S\to R}$ | $hr^{-1}$ | 0 | 1 | 0.195506 | (6) |
| Transport conductance from stem to pod | $\sigma_{S\to P}$ | $hr^{-1}$ | 0 | 5 | 2.527349 | (6) |
| **Senescence Parameters** | | | | | | |
| Max senescence fraction rate (L) | $v_{sene,\max,L}$ | $hr^{-1}$ | 0 | 0.1 | 0.032867 | (8) |
| Max senescence fraction rate (S) | $v_{sene,\max,S}$ | $hr^{-1}$ | 0 | 0.1 | 0.040548 | (8) |
| Max senescence fraction rate (R) | $v_{sene,\max,R}$ | $hr^{-1}$ | 0 | 0.01 | 0.002179 | (8) |
| Senescence transition steepness with DVI (L) | $\alpha_L$ | – | 0 | 10 | 9.190201 | (8) |
| Senescence transition steepness with DVI (S) | $\alpha_S$ | – | 0 | 10 | 8.423220 | (8) |
| Senescence transition steepness with DVI (R) | $\alpha_R$ | – | 0 | 2 | 0.184008 | (8) |
| Senescence midpoint DVI (L) | $\beta_L$ | DVI | 1.5 | 2.5 | 1.815131 | (8) |
| Senescence midpoint DVI (S) | $\beta_S$ | DVI | 1.5 | 2.5 | 2.286124 | (8) |
| Senescence midpoint DVI (R) | $\beta_R$ | DVI | 1.5 | 2.5 | 2.446758 | (8) |
| Retained substrate fraction (LSR) | $f_r$ | – | 0 | 1 | 0.012971 | (10)(11) |
| **DVI switches** | | | | | | |
| DVI threshold for start of pod growth | $\mathrm{DVI}_{\text{pod start}}$ | – | 1 | 1.3 | 1.198307 | – |
| DVI threshold for end of crop growth | $\mathrm{DVI}_{\text{stop growth}}$ | – | 1.8 | 2.2 | 1.920997 | – |

Table S2: Variables in the UTR model

| Variable | Symbol | Unit | Equation |
|---|---|---|---|
| Mass fraction | $MF_O$ | dimensionless | $MF_O = C_{sub,O}/C_{str,O}$ |
| Utilization rate | $r_{u,O}$ | mol m$^{-2}$ hr $^{-1}$ | 3 |
| Respiration rate | $r_{res,O}$ | mol m$^{-2}$ hr $^{-1}$ | 4 |
| Growth rate | $r_{grow,O}$ | mol m$^{-2}$ hr $^{-1}$ | 5 |
| Transport rate | $r_{trans,O_i \to O_j}$ | mol m$^{-2}$ hr $^{-1}$ | 6 |
| Senescence rate of structural C | $v_{sene,str,O}$ | mol m$^{-2}$ hr $^{-1}$ | 9 |
| Senescence rate of substrate C | $v_{sene,sub,O}$ | mol m$^{-2}$ hr $^{-1}$ | 11 |
| Retained substrate C from senesced organ | $r_{reuse,O}$ | mol m$^{-2}$ hr $^{-1}$ | 10 |
| Canopy assimilation rate | $A_{net}$ | mol m$^{-2}$ hr $^{-1}$ | from BioCro |
| Substrate carbon in the organ | $C_{sub,O}$ | mol m$^{-2}$ ground area | 13 |
| Structural carbon in the organ | $C_{str,O}$ | mol m$^{-2}$ ground area | 14 |
| Dry biomass of the organ | $M_O$ | Mg ha$^{-1}$ | 15 |

Table S3: Updated parameters for LD11-2170 vs Pioneer 93B15

| Variable meaning | Variable name | Unit | Pioneer | LD11 |
|---|---|---|---|---|
| Ball-Berry equation intercept | $b_0$ | mol m$^{-2}$ s$^{-1}$ | 0.008 | 0.134 |
| Ball-Berry equation slope | $b_1$ | dimensionless | 10.6 | 5.29 |
| Maximum carboxylation velocity | $V_{cmax}$ | $\mu$mol m$^{-2}$ s$^{-1}$ | 110 | 117.49 |
| Light-saturated potential electron transport rate | $J_{max}$ | $\mu$mol m$^{-2}$ s$^{-1}$ | 195 | 215.07 |
| Max triose phosphate utilization rate | TPU$_{max}$ | $\mu$mol m$^{-2}$ s$^{-1}$ | 13 | 14.48 |
| Electrons per carboxylation | — | dimensionless | 4.5 | 4 |
| Electrons per oxygenation | — | dimensionless | 5.25 | 4 |
| Specific leaf area | iSp | ha Mg$^{-1}$ | 3.5 | 3 |
| Atmospheric $CO_2$ level | $C_{atm}$ | ppm | 372.59 | 414.71 |

Table S4: Leaf starch, sugar (glucose + fructose + sucrose, GFS), and total nonstructural carbohydrate (TNC) concentrations of LD11-2170 sampled in 2022. Starch% and GFS% are the fractions of TNC; SD is the is the standard deviation of TNC.

| | | Starch (mg/g) | | | GFS (mg/g) | | | TNC (mg/g) | |
|---|---|---|---|---|---|---|---|---|---|
| Date (DOY) | Hour | Mean | SD | % | Mean | SD | % | Mean | SD |
| 07/05 (186) | 10:00 | 54.3 | 0.9 | 55.7 | 43.2 | 59.8 | 44.3 | 97.5 | 1.4 |
| 07/05 (186) | 13:00 | 68.8 | 24.7 | 72.5 | 26.1 | 5.3 | 27.5 | 94.9 | 23.3 |
| 07/05 (186) | 16:00 | 165.5 | 1.3 | 85.5 | 28.0 | 38.6 | 14.5 | 193.5 | 8.9 |
| 07/05 (186) | 19:00 | 158.5 | 12.8 | 88.4 | 20.8 | 9.7 | 11.6 | 179.3 | 12.7 |
| 07/05 (186) | 22:00 | 72.2 | 24.9 | 78.5 | 19.8 | 6.8 | 21.5 | 91.9 | 22.5 |
| 07/06 (187) | 01:00 | 32.1 | 22.2 | 60.0 | 21.4 | 2.8 | 40.0 | 53.5 | 21.6 |
| 07/06 (187) | 04:00 | 25.9 | 5.6 | 52.3 | 23.6 | 22.2 | 47.7 | 49.5 | 7.2 |
| 07/06 (187) | 07:00 | 6.1 | 2.0 | 22.8 | 20.6 | 22.8 | 77.2 | 26.7 | 1.8 |
| 07/28 (209) | 07:00 | 22.3 | 5.0 | 39.2 | 34.6 | 42.1 | 60.8 | 56.9 | 0.1 |
| 07/28 (209) | 13:00 | 106.8 | 52.0 | 72.4 | 40.7 | 14.7 | 27.6 | 147.5 | 53.8 |
| 08/24 (236) | 07:00 | 129.3 | 50.5 | 89.5 | 15.2 | 43.2 | 10.5 | 144.5 | 50.8 |
| 08/24 (236) | 13:00 | 195.1 | 17.1 | 92.7 | 15.4 | 3.8 | 7.3 | 210.6 | 15.2 |
| 08/24 (236) | 19:00 | 228.2 | 43.7 | 94.1 | 14.2 | 34.0 | 5.9 | 242.4 | 44.1 |
| 09/15 (258) | 07:00 | 34.4 | 10.9 | 74.8 | 11.6 | 1.2 | 25.2 | 46.0 | 10.9 |
| 09/15 (258) | 13:00 | 22.0 | 13.7 | 62.7 | 13.1 | 1.7 | 37.3 | 35.1 | 12.2 |
| 09/15 (258) | 19:00 | 14.0 | 12.6 | 54.5 | 11.7 | 1.7 | 45.5 | 25.6 | 12.1 |

Table S5: Stem starch, sugar (glucose + fructose + sucrose, GFS), and total nonstructural carbohydrate (TNC) concentrations of LD11-2170 stems sampled in 2022.

| | | Starch (mg/g) | | | GFS (mg/g) | | | TNC (mg/g) | |
|---|---|---|---|---|---|---|---|---|---|
| Date (DOY) | Hour | Mean | SD | % | Mean | SD | % | Mean | SD |
| 07/05 (186) | 10:00 | 39.2 | 54.0 | 36.2 | 68.9 | 22.6 | 63.8 | 108.1 | 66.9 |
| 07/05 (186) | 13:00 | 23.6 | 13.8 | 27.6 | 61.9 | 73.4 | 72.4 | 85.5 | 41.3 |
| 07/05 (186) | 16:00 | 24.0 | 14.5 | 32.7 | 49.3 | 46.1 | 67.3 | 73.2 | 18.0 |
| 07/05 (186) | 19:00 | 23.4 | 15.0 | 37.4 | 39.2 | 30.0 | 62.6 | 62.6 | 18.9 |
| 07/05 (186) | 22:00 | 59.2 | 15.3 | 65.5 | 31.2 | 26.3 | 34.5 | 90.3 | 7.5 |
| 07/06 (187) | 01:00 | 52.3 | 42.2 | 51.3 | 49.7 | 14.7 | 48.7 | 102.0 | 49.2 |
| 07/06 (187) | 04:00 | 10.0 | 12.5 | 15.7 | 53.7 | 60.2 | 84.3 | 63.6 | 44.9 |
| 07/06 (187) | 07:00 | 9.4 | 5.9 | 30.8 | 21.1 | 20.6 | 69.2 | 30.5 | 14.9 |
| 07/28 (209) | 07:00 | 9.0 | 7.7 | 6.1 | 139.3 | 169.3 | 93.9 | 148.3 | 80.6 |
| 07/28 (209) | 13:00 | 13.9 | 5.5 | 10.6 | 117.6 | 135.5 | 89.4 | 131.5 | 42.3 |
| 08/24 (236) | 07:00 | 19.3 | 4.4 | 32.2 | 40.5 | 45.6 | 67.8 | 59.7 | 9.2 |
| 08/24 (236) | 13:00 | 21.7 | 11.2 | 32.1 | 45.9 | 44.1 | 67.9 | 67.6 | 2.5 |
| 08/24 (236) | 19:00 | 34.6 | 22.2 | 52.0 | 32.0 | 20.7 | 48.0 | 66.5 | 23.6 |
| 09/15 (258) | 07:00 | 4.1 | 3.9 | 6.1 | 63.2 | 83.8 | 93.9 | 67.3 | 40.6 |
| 09/15 (258) | 13:00 | 26.6 | 36.2 | 24.2 | 83.4 | 89.4 | 75.8 | 110.0 | 23.3 |
| 09/15 (258) | 19:00 | 8.9 | 6.5 | 10.8 | 73.3 | 97.0 | 89.2 | 82.2 | 15.2 |
| 10/05 (278) | 12:00 | 0.0 | 0.0 | 0.0 | 2.6 | 2.6 | 100.0 | 2.6 | 0.0 |

Table S6: Reproductive stages with corresponding vegetative stages and first observation dates for LD11-2170 in 2023 and 2024. Some reproductive stages were not recorded in the survey. Their corresponding vegetative stages and first observed dates are shown as "–".

| Reproductive Stage | Vegetative stages | | First observed date | |
|---|---|---|---|---|
| | 2023 | 2024 | 2023 | 2024 |
| R1 | V7 | V8–V10 | 7/21 | 7/12 |
| R2 | V8–V10 | V10–V11 | 7/26 | 7/19 |
| R3 | – | V10–V12 | – | 7/29 |
| R4 | V13 | – | 8/08 | – |
| R5 | V13 | V13–V16 | 8/16 | 8/07 |
| R6 | V14–V15 | V15–V16 | 8/24 | 8/21 |
| R7 | V15 | – | 9/26 | – |
| R8 | V15 | – | 9/29 | – |
